# Shape-shifting microgel automata controlled by DNA sequence instructions

**DOI:** 10.1101/2022.09.21.508918

**Authors:** Ruohong Shi, Kuan-Lin Chen, Joshua Fern, Siming Deng, Yixin Liu, Dominic Scalise, Qi Huang, Noah J. Cowan, David H. Gracias, Rebecca Schulman

**Author notes:** Voiland School of Chemical Engineering and Bioengineering, Washington State University; Pullman, WA 99164, USA.

## Abstract

Controlling material shapes using information-bearing molecular signals is central to the creation of autonomous, reconfigurable soft devices. While physical and chemical stimuli can direct simple material swelling, bending, or folding, it has been challenging to direct multi-step shape-change programs crucial for complex, robotic tasks. Here, we demonstrate gel automata— sub-millimeter, photopatterned, highly swellable DNA gels—whose parts grow or shrink in response to easily designed DNA activator sequences, allowing for precisely controlled device articulation. We design and fabricate gel automata that reversibly transform between different letter shapes, and use neural networks to design automata that transform into every even or every odd numeral via designed reconfiguration programs. This sequential and repetitive metamorphosis of materials via chemical reorganization could dramatically advance our ability to manipulate micro-particles, cells, and tissues.

**One-Sentence Summary:** Photopatterned microgels follow sequences of DNA instructions to transform between complex, meaningful shapes such as letters and numerals.

## Main Text

An automaton is a machine that can repeatedly take on different configurations in response to corresponding instructions. The ability to specify a sequence of these instructions, *i*.*e*., a program, underlies the astonishing power of automata, which can keep time, write, or play instruments (*1–3*). Modern robots usually interpret electronic programs, but automata that interpret mechanical instructions date back to Archytas of Tarentum’s wooden dove (*4, 5*).

Nucleic acids and proteins can act as biochemical instructions that direct complex and critical biological processes such as morphogenesis, growth, and disease fighting (*6, 7*). Biomolecules have unique advantages as signals: they can be produced in astonishing variety, diffuse through three-dimensional media, and react with high specificity (*8*). Biomolecules can provide energy to drive chemical reactions and can also become structural components of material (*9, 10*). In contrast, light or electronic signals require batteries or external power sources to store energy and line-of-sight access, wires, or tubes that direct signals to a specific location, where they interact with limited specificity (*11–13*).

We asked whether we might develop a means to use biomolecular signals to program different, repeated shape changes of synthetic soft micro-devices. As in biological tissues, localized growth or shrinkage could produce precise changes in the curvature of different regions, which might also be tuned to control global shape change (*14–16*). Multiple such deformations performed in sequence could reversibly transform a material between many different global configurations.

Here, we (1) design specific and selective DNA instructions for shape change and the complementary gels that respond to them, (2) develop scalable methods for fabricating multi-segmented gels responsive to these different instructions, and (3) engineer hydrogel curvature and mechanical properties so that gel automata execute complex shape-change programs encoded as DNA sequences.

We use gels composed of polyacrylamide/bis-acrylamide (PAAM-co-BIS) or polyethylene glycol (PEG) polymer backbones that are cross-linked by DNA. The hydrogel shape change is driven by DNA hybridization-induced polymerization (*17–19*). DNA strands are inexpensive, routine to produce, easy to store, and generally biocompatible (*20–22*). A plethora of DNA circuits and responsive release processes could produce DNA sequences to drive a series of such interactions and induce adaptive responses without human intervention (*23, 24*).

Shape change of ultrathin films can be directed using DNA signals (*25–27*). DNA sequences can also swell or shrink hydrogels (*17, 18, 28–31*), but afterward, these hydrogels are no longer responsive to DNA instructions. To direct multi-step gel growth or shrinking programs, we first sought to design a mechanism for growing a hydrogel using one DNA sequence instruction and then shrinking the hydrogel using a second DNA sequence instruction, thereby reversing the growth process. We investigated a variety of processes to shrink PAAM-co-BIS-DNA and PEG-co-DNA gels selectively. One involved a zipping and unzipping mechanism, in which one pair of DNA sequences induces growth by processively polymerizing at DNA crosslinks in a reversible but forward-driven process, and another pair causes shrinking by altering the polymerizing monomers’ conformations, so processive depolymerization becomes favored (fig. S1). However, such a process failed to induce gel shrinking (fig. S2), possibly due to sluggish kinetics (Supplementary Text S1).

Two other experiments established that the depletion of DNA from a gel could drive its shrinking. In the first experiment, we grew PEG-co-DNA gels by adding growth activators, then treated the gels with DNase I, an enzyme that cleaves DNA’s phosphodiester linkages (fig. S3). The gels shrunk back to their original sizes. However, because DNase I irreversibly cleaves the crosslinks that serve as initiation points for growth activator polymerization, adding more activators did not initiate the second growth cycle. In the second experiment, we expanded gels using growth activators, then denatured them by heating them to 90ºC. This process also caused gels to shrink back to their original sizes (fig. S4), presumably because the heat de-hybridized the polymerized DNA crosslinks. De-hybridization is reversible, so the crosslinks might reform after cooling. Indeed, the gels could be re-grown again with the addition of growth activators after cooling. However, heat acts upon the entire gel and does not offer the capacity to shrink particular regions of a gel selectively. In short, it did not provide the level of programmability that can be achieved by DNA-directed gel shrinking.

We built on the idea that rapid crosslink depolymerization induced gel shrinking to design two sets of DNA sequences that direct growth and shrinking, respectively. One pair of hairpin-shaped growth activators processively polymerize at crosslinks to expand a specific gel region. Another pair of shrinking activators bind to and sequester specific growth activator strands, denaturing the DNA polymers and breaking the crosslinks to shrink a particular gel region (Fig. 1A). An initial pair of growth/shrinking activators (Table S1, SDE_v1) induced modest growth then shrinking of a corresponding hydrogel. Guided by the thermodynamics of DNA-crosslinked gel shape change and the kinetics of toehold-mediated 4-way DNA branch migration (*17–19, 32*), we varied the domain lengths of these DNA activators to find designs that maximized the amount of growth and shrinking observed (Fig. 1C, fig. S5). Increasing the relative concentrations of the optimized growth/shrinking activators then further increased the extents of growth and shrinking (Fig. 1D). We also found that combining growth activators with a small concentration of DNA sequences that terminated extension further increased the amount of growth observed (fig. S6), perhaps because these “terminator” sequences limited the length of polymers that formed in the absence of an initiator, preserving growth fuel for actuation. Using these results, we designed four systems of growth/shrinking activator sequences that could each specifically and repeatedly direct the growth and shrinking of both PEG and PAAM-co-BIS DNA gels with corresponding crosslink sequences (Fig. 1E-F, fig. S7-8). DNA sequences that were not purified after synthesis and thus were inexpensive reliably grew PAAM-co-BIS-DNA and PEG-co-DNA gels as much as 10 and 4 times volumetrically and then shrank over five cycles (fig. S9).

**Fig. 1.**
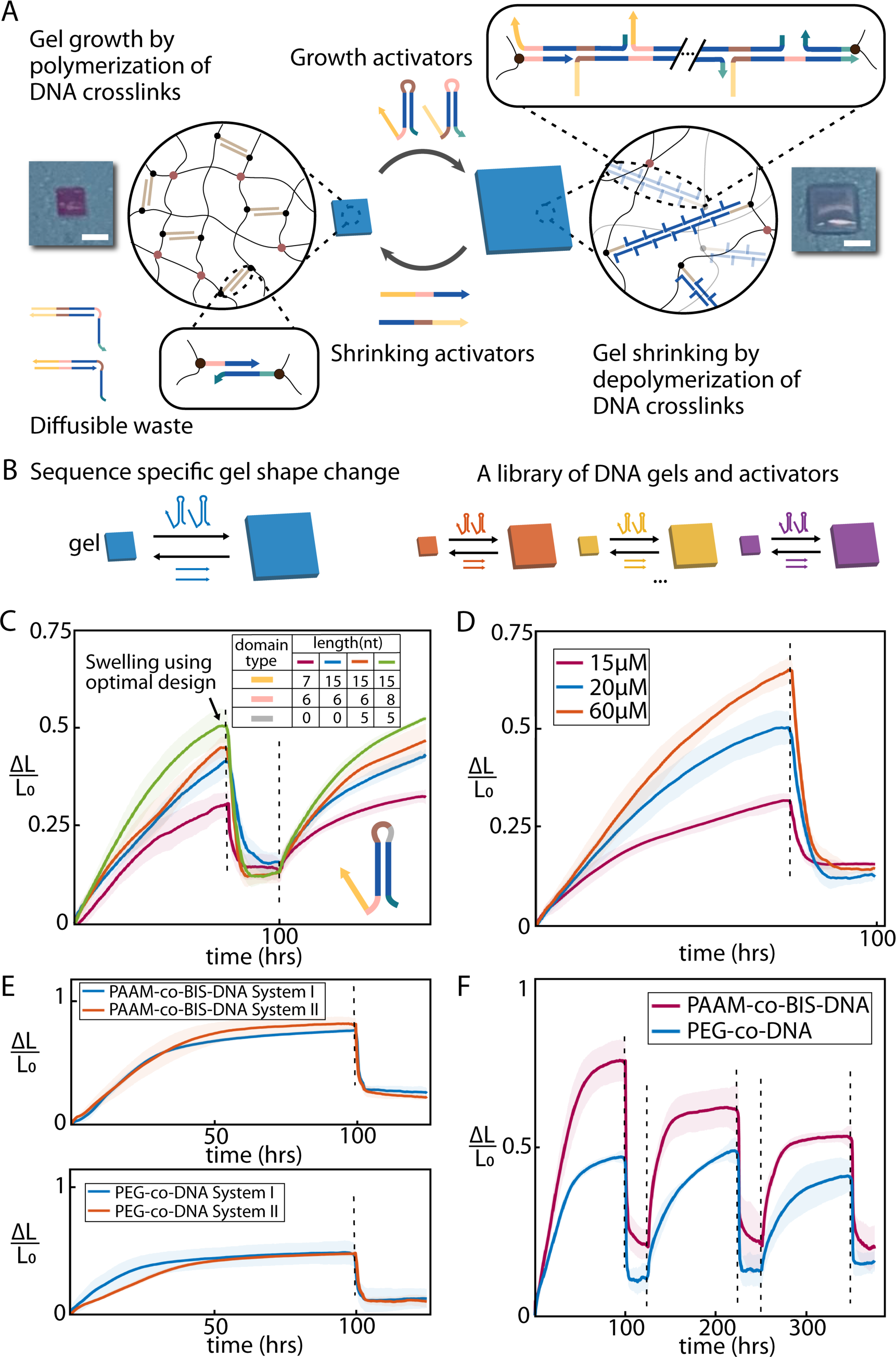
Tunable DNA gel growing and shrinking directed by DNA activators through strand displacement. **(A)** Schematic of DNA-directed gel growth and shrinkage. Hairpin-shaped growth activators activate the growth of DNA gels, while the shrinking activators direct the reverse reaction. Black dots, Acrydite-DNA linkers. Brown dots, covalent chemical crosslinks. Scale bar, 1 mm. **(B)** Colors of the DNA gel represent different sequences of DNA crosslinks (left to right, Table S1: System I, II, III, IV A and R strands) co-polymerized with gel polymer backbone. Corresponding activators induce orthogonal growth and shrinkage. **(C)** Parameter study of activator designs via varying overhang domain (yellow), primary toehold (pink), and HP spacer (grey) lengths. Table S1: Sequence Design Evolution strands. PEG-co-DNA gels grew, shrank (solution changes indicated by dotted lines), and re-grew. Solid lines represent the average change in the side length of the gel with respect to the original side length (N = 3). Shaded areas represent a single standard deviation from the mean growth value. **(D)** Parameter study of gel actuation with varying [activators]. The System I PEG-co-DNA gels were treated with different [activators], protocol as in Fig. 1C. **(E)** Kinetics study of PAAM-co-BIS-DNA and PEG-co-DNA gels. System I and II gels were treated with [activators] = 60 µM with 1% terminator strands, other steps as in Fig. 1C. **(F)** Three actuation cycles of System I gels, protocol as in Fig. 1E.

We next sought to explore whether selective growth and shrinking of different gel regions could deform multi-segmented gel automata. We photopatterned gel bilayers and developed instructions for actuation primitives such as curve, uncurve, expand, and invert, directed by specific actuation sequences that could be combined into programs (Fig. 2A). We ran the resulting shape-change programs to characterize how one or both layers of bilayers grew, shrank, and induced curvature. The ability to use orthogonal DNA signals for growth and shrinking made it possible to simultaneously grow some gel regions while shrinking others, offering unprecedented control over pathways of shape change. Forces generated by both gel growth and shrinking could drive shape changes such as bending, straightening, turning, and lengthening (Fig. 2B-D, Movies S2-4). Our system also exhibited control over curvature extent; bilayers could curve slightly or at least into a circle (1.2-2 mm^-1^ curvature, fig. S10), and, because shrinking proceeded faster than growth, this allows control over relative curving rate. Growth and shrinking programs could direct reversible deformation over at least three cycles (Fig. 2E, Movie S5).

**Fig. 2.**
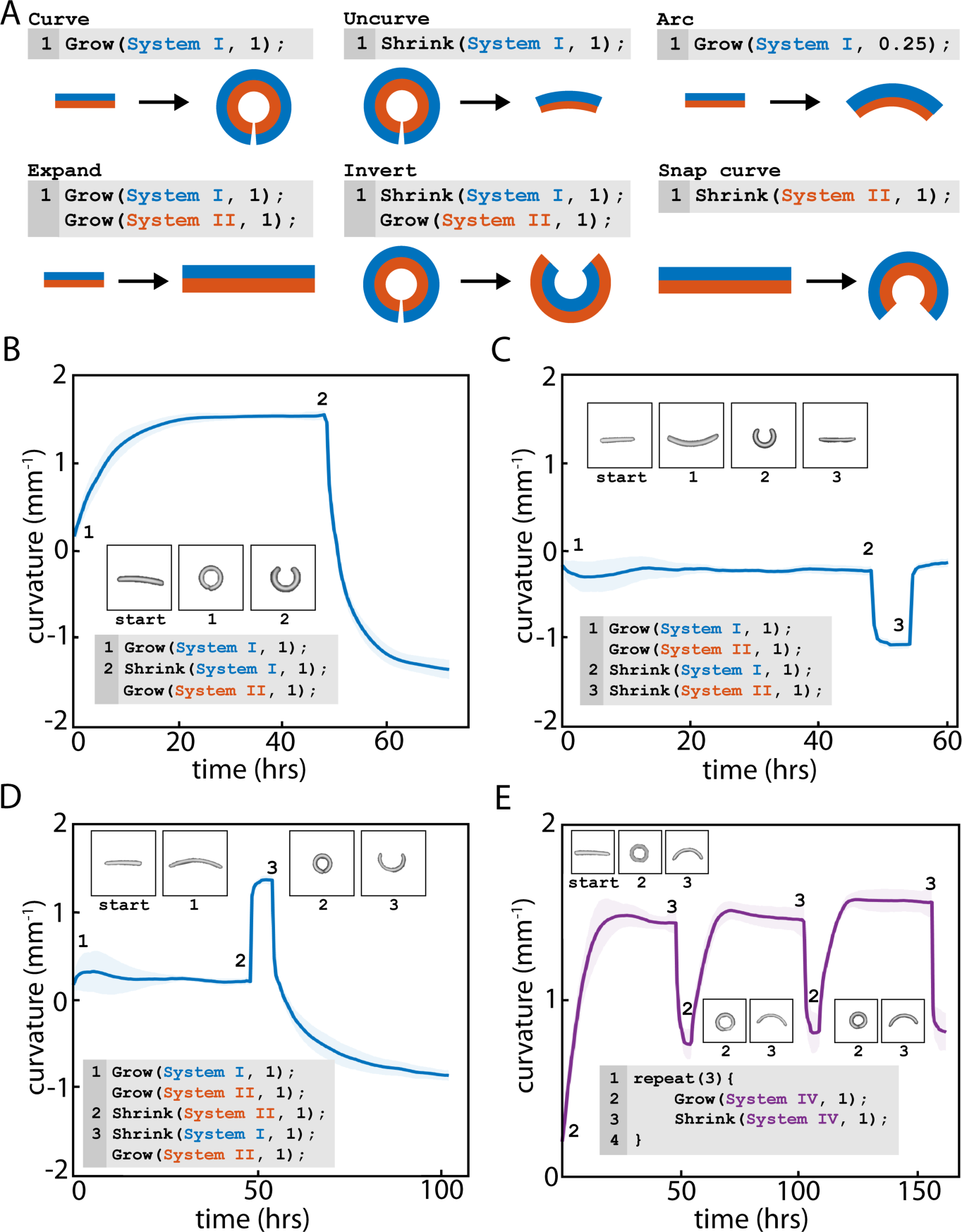
Programs for gel bilayer shape change by selectively growing and shrinking specific gel regions. **(A)** Example behaviors (curve, uncurve, etc.) and corresponding programs that instantiate each gel bilayer transformation. We define each actuation instruction using the syntax action(system, degree). The first argument identifies the target DNA system, and the second argument specifies the degree of shape change (1 being the maximum), which can be specified using the concentration of actuators supplied (for 1, 60 µM, and 0.25, 15 µM). The example gels consist of a System I DNA gel layer (blue) and a System II DNA gel layer (red). **(B-D)** Curvature changes of PEG-co-DNA System I-System II bilayer gels during different actuation programs. Each statement was executed at the time indicated by the numbers near each plotted line. Gel micrographs with numbers show the steady state gel shapes after executing the corresponding statement. Gel size: 3.2 mm x 0.8 mm x 0.2 mm at the start of actuation. For protocol details, see Materials and Methods. **(E)** Curvature changes of PEG-co-DNA System III-System IV bilayer gels during the program’s execution; other details are the same as described in Fig. 2B-D.

This advanced and flexible control over the degree of actuation, speed, and curvature direction suggested that gel automata programs could direct complex, multi-step bending and shape transformation processes. To test this hypothesis, we asked whether we could design multi-segment gels where the growth and shrinking of specific gel regions could allow the gels to transition between many distinct functional shapes. We first sought to design gel automaton strips that could present **I**-, **J**-, **S**-, and **C**-shaped curves. We calculated the segment lengths of bilayer strips that would transform between these four shapes and designed an actuation program to switch between them (Fig. 3). Fabricating this structure required four computer-aided design (CAD) masks and four photopatterning alignment steps. We modified a previously developed photolithographic gel patterning protocol (*17*) to reduce misalignment, form gel regions with uniform thicknesses, reduce delamination from the substrate during patterning, and allow for reliable lift-off of complete, intact devices. Photomasks were used for exposure and as the underlying substrate in multiple patterning steps to improve alignment and reduce material waste. DNA gels are optically transparent; co-polymerizing adjacent gel regions with different dyes allowed for the precise visual alignment required to fabricate complex, multi-segmented gels. We also created CAD masks for adjacent gel segments with overlaps and optimized overlap size to balance material usage, binding firmness, and alignment accuracy (Supplementary Text S2, fig. S11).

**Fig. 3.**
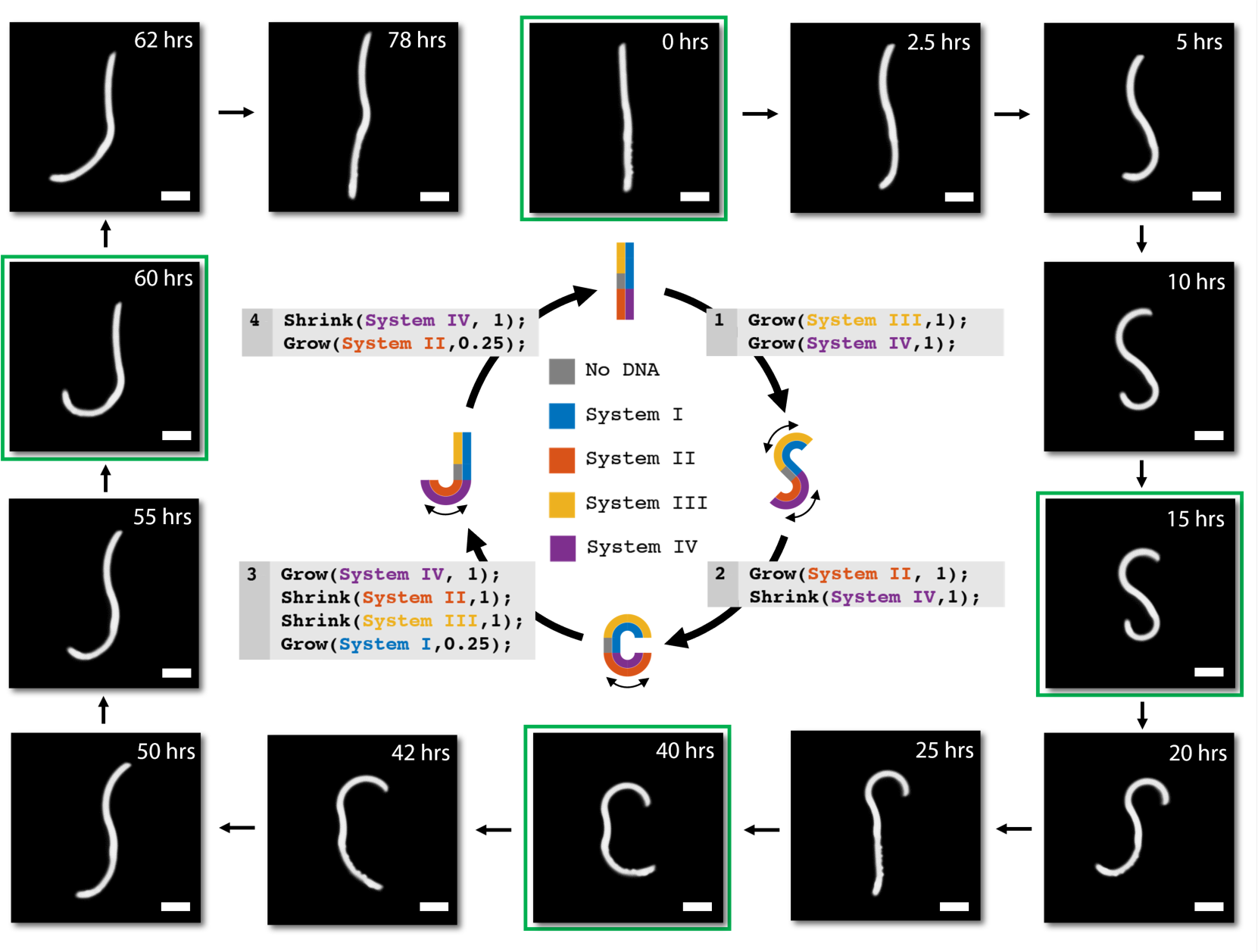
Letter gel automaton design and DNA program guiding actuation. Design (inner part of the figure) of a letter gel automaton that transforms into **I, S, C**, and **J** shapes and time-lapse fluorescence micrographs (outer part of the figure) of the letter gel during program execution. The micrographs in the green boxes show the gel automaton’s state right before the program executes each step (1-4). Curved arrows around gel automaton schematics indicate gel regions being directed to grow in the current step. Design parameters, actuation details, and actuation video are in Supplementary Text S2, fig. S11, S13, and Movie S6. Scale bars, 1.5 mm.

During transformation tests of an initial design (V1.0, fig. S12) for the letter gel automaton, the letter **I** successfully transformed into an **S** and a **C**. However, the top part of the automaton did not fully straighten during transformation to a **J**. The top part remained curved as the automaton returned to the **I** configuration. Hypothesizing that this errant curvature was due to one region remaining slightly more swollen after growth and shrinking than it was in its original state, we revised the automaton’s design. In letter automaton V1.1, we replaced one of the regions without DNA crosslinks in letter automaton V1.0 with a System I gel (fig. S13). Furthermore, we altered the transformation program for entering the **I** state from **J** to include an instruction that slightly grew this region to straighten the segment that had remained curved. Letter automaton V1.1 then transformed between **I, S, C**, and **J** shapes and returned to its original form as directed by DNA signals (Fig. 3, Movie S6).

The use of four distinct binary DNA systems allows for up to 2^4^ =16 actuation states. Altering the curvatures of different segments in a multi-segment structure could specify a range of complex forms. To demonstrate such programmable and non-trivial shape change, we sought to develop gel automata that could transform into different Arabic numerals, which might be useful for communicating numeric messages. In the **I**/**S**/**C**/**J** automaton, two specific segments were modularly designed to curve and un-curve into the top and bottom shapes of each of the letters. In contrast, forming the diverse shapes of Arabic numerals would require different segments to play different roles when forming different numerals. As a result, there was no obvious way to break the design problem into pieces. We hypothesized that we could design such structures in a manner reminiscent of designing biomolecular sequences (*33*): develop an automated algorithm to predict the configurations of different potential “digit automaton” device designs for each of the 16 simulated actuation states, and then search through design space for structures that properly formed the desired set of digits.

To create a simple simulator for strip bending that could be used for design, we measured the curvatures of bilayer gels made from different combinations of DNA system crosslinks (fig. S10) and assumed that each segment of a multi-segment structure would achieve the same curvatures as they would in isolated bilayers. This simulator predicted the shapes of candidate devices in each of the specific actuation states and produced simulated micrographs. We then trained a convolutional neural network (CNN) to classify these simulated images to evaluate how well candidate designs could be recognized visually as different digits. Then, we developed a genetic algorithm to evolve designs for devices that could form the set of odd digits or the set of even digits (Fig. 4A, fig. S14-15). To ensure our designs could be fabricated reliably, we restricted our search to devices that could be fabricated in five or fewer steps using the fabrication process we developed. We were able to design and then fabricate one even digit and one odd digit automaton design (Fig. 4B, fig. S16-17). These structures were flat in their initial configuration as designed, indicating precise fabrication, alignment, and lift-off over the multiple required fabrication steps (fig. S18). When we actuated each automaton using a program that switched between the actuation states, the resulting structures transformed into each of the designed digit shapes in a prescribed sequence, either **1, 3, 5, 9, 7** for the odd digits (Fig. 4C), or **1** (the initial flat strip), **0, 2, 6, 8, 4** for the even digits (Fig. 4D). Extra DNA activators were added at a couple of later stages to increase/decrease specific curvatures as stated in Fig. 4B. While mechanical stresses generally cause long-range interactions between adjacent elastic columns (*34*), the assumption that the curvature of different segments could be viewed modularly led to remarkably good predictions of shapes: the digits formed by the automata were recognizable and almost identical to the predicted shapes. The availability of different activators also allowed a high degree of flexibility in precisely adjusting the sizes of targeted regions of the gel devices to achieve the desired final shapes.

**Fig. 4.**
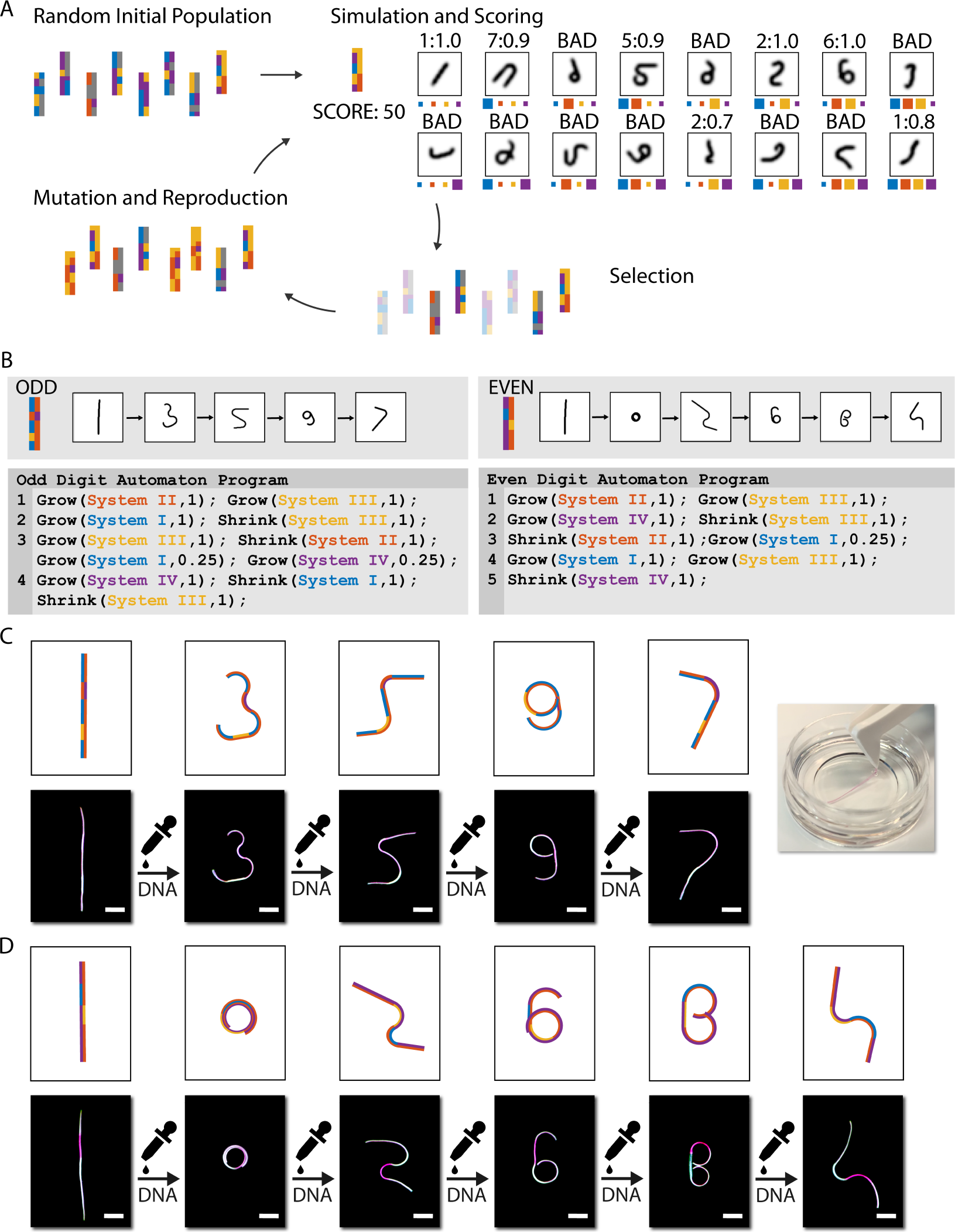
Neural network-guided design of multistate digit gel automata. **(A)** Schematics of the machine learning model simulation, scoring, mutation, and selection process. Colors of the gel regions represent different systems of DNA crosslinks in a gel, as in Fig. 3. **(B)** Optimized even and odd digit gel automata designs and actuation programs. **(C) (D)** Actuation of digit gel automata. Odd **(C)** and even **(D)** digit automata were actuated with respective DNA activation solutions following the programs in Fig. 4B. DNA gel regions were labeled with either fluorescein-O-methacrylate (white) or methacryloxyethyl thiocarbamoyl rhodamine B (magenta) during fabrication for easier alignment; these colors do not correspond to the colors in the schematics. Top right, a photograph of an odd gel automaton. Scale bars, 5 mm.

In summary, we have demonstrated mass-producible, shape-shifting gel automata in which biomolecules dynamically and reversibly trigger the manipulation of architected material shape at the molecular scale. Automata structures and their complex transformation protocols can be expressed, respectively, as mask designs and actuation programs, allowing the enumeration of a wide array of dynamic, multi-functional, and autonomous devices that can be designed using automated workflows and fabricated using wafer-scale methods. The vast space of DNA sequence design allows extensive tuning and programming of complex shape changes, providing a pathway to creating digital materials. It is noteworthy that biochemical activators and repressors direct most signaling and actuation programs in biological systems; their use here transcends the limits of pneumatic, fluidic, or electronic soft activators by allowing ready miniaturization, precise control of shape change, and actuation at a distance and in confined and tortuous spaces. These observations suggest how biochemically controlled autonomous soft robots could intelligently perform complex tasks in various aqueous, marine, and biological environments.

## Author contributions

RSh, JF, DHG, and RSc conceptualized the study. RSh fabricated gels, actuated bilayer gel and gel automata, curated and analyzed data into figures, and created videos under the supervision of DHG and RSc. RSh, KLC, JF, and YL performed monolayer gel actuation parameter studies under the supervision of RSc. RSh, KLC and SD built a programmable imaging system, performed image processing, and designed gel automata under the supervision of NJC and RSc. RSh and KLC designed schematics in the figures. KLC and RSc designed a digit simulator and trained the CNN with input from RSh, NJC, and DHG. DS and QH investigated the processive shrinking directive. RSh, KLC, DHG, and RSc wrote the manuscript with input and edits from all authors.

## Supplementary Materials

Materials and Methods

Supplementary Text S1 to S3

Figs. S1 to S19

Tables S1

Captions for Movies S1 to S6

Movies S1 to S6

