## Supporting Information for "Shape-shifting microgel automata controlled by DNA sequence instructions"

**This PDF file includes:**

Materials and Methods
Supplementary Text S1 to S3
Figs. S1 to S19
Tables S1
Captions for Movies S1 to S6

**Other Supplementary Materials for this manuscript include the following:**

Movies S1 to S6

### Materials and Methods

**DNA Sequences and Preparation**. The sequences for the DNA crosslinks and growth and shrinking activators are listed in Supplementary Materials Table S1. DNA Sequences were designed using NUPACK 3.2.2 to have specific secondary structures and minimal undesired crosstalk (*35*). A temperature of 25°C and salt conditions of 0.05 M $Na^+$ and 0.0125 M $Mg^{2+}$ were used in all design programs. Designs were produced iteratively by adding new sequences to an existing set of sequences and domains (system 1-4 strands in (*17*)); multiple design trials were run to produce several potential sets of sequences. These sets of sequences were then ranked by the degrees of interaction with existing sequences predicted by NUPACK for the final selection of sequence designs. Sample scripts are at https://github.com/charliecharlie29/Gel-Automata-Directed-By-DNA-Codes. Unmodified and Acrydite-modified oligonucleotides were purchased in lyophilized form from Integrated DNA Technologies (IDT) with standard desalting purification. The DNA strands were solubilized in Tris-acetate-EDTA (TAE) /0.0125 M $Mg^{2+}$ (TAEM) buffer (TAE buffer, Life Technologies, #24710-030; Magnesium acetate tetrahydrate, Sigma #228648). The DNA's concentration in solutions was verified using absorbance spectroscopy at 260 nm. 3 mM DNA crosslink complexes were annealed in TAEM from 90°C to 20°C at a rate of 1°C/min using an Eppendorf Mastercycler. Growth activator strands were heated to 95°C for 15 mins and then flash-cooled in ice for 5 mins at a concentration of 400 μM.

**Preparation of DNA Gel Pre-gel Solution.** Solutions of hydrogel monomers and DNA crosslinks were prepared as reported previously by (*18*). Briefly, the concentrations of the components in PAAM-co-BIS-DNA pre-gel solution are: 1.41 M of acrylamide (BIO-RAD #161-0100), 5 mM of N, N'-methylenebis(acrylamide) (Sigma-Aldrich, #146072), 1.154 mM DNA crosslinks, 2%v/v Omnirad 2100 (iGM Resins USA, #55924582), and 2.74 mM methacryloxyethyl thiocarbamoyl rhodamine B (Polysciences, Inc., #23591). The concentrations of the components in the PEG-co-DNA pre-gel solution are the same as those in the PAAM-co-BIS-DNA pre-gel solution except PEGDA-MW10k (Sigma-Aldrich, #729094) and PEGDA-MW20k (Sigma-Aldrich, #767549) were 10wt%, and one of these was used in place of acrylamide and bis-acrylamide. Omnirad 2100 was first made into a 75%v/v butanol solution to help disperse into the pre-gel solution. When making gel bilayers, 1 mM fluorescein-O-methacrylate (Sigma, #568864) fluorescent dye was included in lieu of 2.74 mM methacryloxyethyl thiocarbamoyl rhodamine B (Polysciences Inc., #23591) in the pre-gel solution when patterning the second (upper) gel layer. The pre-gel solutions were mixed well using a pipettor and then were ultrasonically mixed for 1 min (for PAAM-co-BIS-DNA) or 3 mins (for PEG-co-DNA) before being degassed in a vacuum chamber for 15 mins.

**Lithography Chamber Fabrication.** The photolithography chambers were assembled following a previously reported protocol (*17*, *18*). The chamber consisted of a chromium (Cr) mask, a glass substrate (or a Cr-coated glass substrate with patterns), and desired thickness tape serving as spacers (160 μm for monolayer gels and 60 μm for bilayer gels) to control gel thickness. Plastic masks, which were then utilized for making Cr masks or Cr-coated glass substrate, were first designed using AutoCAD and then sent to Fineline Imaging for printing. Glass slides were cleaned with DI water and isopropyl alcohol (IPA), then blow-dried using nitrogen gas before being spin-coated 3 nm SC1827 (Microchem, Microposit S1800 Series) and baked on a 115°C hotplate for 1 min. The prepared glass slides were cured through plastic masks with 180 mJ/$cm^2$ 365 nm UV light, then developed using a 1:5 w/w Microposit 351 Developer (Shipley) and washed with DI water. A 150 nm (for monolayer gel fabrication) or 300 nm (for multi-step patterning process) Cr layer was then deposited onto the glass slides through thermal evaporation. The remaining photoresist layer was washed by acetone ultrasonically and DI water. All glass slides, Cr masks, and Cr-coated glass substrate were cleaned with water and IPA and then blow-dried before each use. For multi-step photopatterning, CYTOP (Type M, Bellex International Corp.) was applied to Cr masks to prevent gels from sticking or lift-off.

**Photopatterning Process.** To pattern square-shaped, 1mm side-length monolayer gel films with a thickness of 160 μm, the pre-gel solution was injected into a chamber and then exposed to a 365 nm UV light source for 160 mJ/cm$^2$ (PAAM-co-BIS), 800 mJ/cm$^2$ (PEGDA-MW10K) or 1000 mJ/cm$^2$ (PEGDA-MW20K). To pattern 2 mm-long, 0.5 mm-wide bilayer gel structures with each layer thickness of 60 μm, the pre-gel solution with fluorescein-O-methacrylate was first patterned using the exposure energy stated above. The first gel layer was gently washed using TAEM buffer and dried using nitrogen gas. The second 60 μm spacer was added to the gel chamber to increase the chamber height to a total of 120 μm. The mask was then aligned with the patterned gel, and the pre-gel solution containing methacryloxyethyl thiocarbamoyl rhodamine B was UV-cured. Process diagrams for the multi-step gel automata photopatterning process are shown in fig. S11, fig. S16, and fig. S17 and additional detailed descriptions of the processes are given in Supplementary Text S2. Briefly, we designed the Cr masks to have four aligning cross-shaped markers at the corners to enable alignment between gel segments. The final segment lengths of each region in the photopatterned masks for the multi-segment strips were each 1.5 times the lengths designed using computer programs. This increase in mask length was introduced to compensate for the curvature reduction in the subsequent cycle of a multi-step actuation. During the first step of fabrication of multi-segment structures, a Cr-coated glass substrate with the same pattern as the photomask was used instead of a transparent glass substrate so that the aligning markers on the Cr mask could be used during subsequent patterning steps. In the following fabrication steps, gels were gently washed with TAEM and blow-dried after each patterning step, and additional spacers were added when the first layer of photopatterning was finished, as described in the bilayer gel fabrication process. All gels were washed and hydrated with TAEM buffer, gently removed from the glass substrate or Cr-coated glass substrate, and stored in TAEM in a 4°C fridge until actuation.

**Characterization of Monolayer DNA Gel Shape Change.** The extent of growth and shrinking of monolayer gels were measured using time-lapse fluorescence imaging with a gel imager (Syngene EF2 G: Box) equipped with a blue light transilluminator (Clare Chemical, max emission at ≈450 nm) and a UV 032 filter (Syngene, bandpass 572–630 nm), or an automated "Pi-Imager" described in the Supplementary Text S3 as "Programmable Imaging System (Pi-Imager) for Time Lapse Fluorescence Image Capturing." The gel samples were transferred to wells within a black-walled 96-well plate to isolate the gels from each other during actuation. Unless noted otherwise, solution composition and solute concentrations are as stated below: gels were expanded in TAEM supplemented with 0.01%v/v Tween20 (Sigma, #051M01811V) (TAEM-Tween20) to prevent gels from sticking to the well surfaces. 150μL of TAEM-Tween20 solution containing 60 μM DNA growth activators (99% polymerizing, 1% terminating) was added to each well. After 72-100 hours of growing, the DNA solution was switched to 100 μL TAEM-Tween20 for 15 mins, and the solution was removed, and then 150 μL TAEM-Tween20 shrinking activators solution was added. The above growth/shrinking process was repeated when characterizing multiple actuation cycles. Images were taken every 30 mins, and the resulting photos were segmented into smaller images, each containing one gel for further MATLAB processing. All images were first transformed into grey-scale images by extracting the red channel signals from the original RGB image. Images were then contrast-stretched using MATLAB's Image Processing Toolbox (2019a) to reduce background and simplify further feature extraction. The four side lengths of the gel in each image were measured and averaged as follows. The extrema and centroids of the objects were determined using MATLAB's function `regionprops`. Eight locations provided by the extrema of a gel object (two points at the ends of each side) were used to determine the locations of the four vertices by K-means clustering. The average distance between these four clusters was used as the measure of the side length of the gel. The relative change in side length ($\Delta L/L_0$) of the gel was calculated using the measured side lengths (L) from each image in a time series relative to the side length prior to adding DNA activators ($L_0$). PAAM-co-BIS-DNA gels were sometimes too dim to find with the MATLAB code described above. In such cases, the raw images were first treated by flat-fielding to have a brighter view and a more significant contrast against the background, in addition to the previously described process for edge-length determination. Shaded area/dotted lines enclosed standard deviation of the mean growth value, which were smoothed using MATLAB `smooth` function over 50 (growing) and 10 (shrinking) points, N (sample size) = 3 or 4.

**Characterization of Bilayer DNA Gel Bending.** Micrographs of bilayer gels were captured using the gel imager, blue light, and filter stated in the "Characterization of Monolayer DNA Gel Shape Change" section in Methods. Before actuation, bilayer gels were set on their sides so that curvature could be measured from a side view of a gel. The DNA-directed actuation and solution exchange processes were the same as those described in the section on monolayer gel shape change characterization. Images were taken every 30 mins and were segmented into images, each containing one gel for further processing in MATLAB. Within each image, the pixels that contained the gel were first determined by selecting pixels more than 4.3 standard deviations brighter than the mean fluorescence intensity of the image to produce a binary image. The binary image was smoothed and thinned to a curve using MATLAB's `bwmorph` function. The radius of curvature of the contour connecting the coordinates on the line/ring was determined using the Taubin method (*36*). The direction of curvature of bilayer gels was distinguished using +/- (the sign of the radius of curvature). Shaded area/dotted lines enclose the standard deviation about the mean curvature value, which were smoothed using MATLAB `smooth` function over 20 (ascending) and 10 (flat and descending) points, N = 3.

**Letter Gel Automata Design.** We began creating a letter gel automaton by manually designing a three-segment gel bilayer stack, with the idea that the top and bottom portions would curve in different directions to form the top and bottom curved elements of the **C**, **S**, and **J**. To choose the lengths and systems of each gel region, we visualized the resulting curves produced by candidate bilayer strips by assuming that the segment of the bilayers would have a radius of curvature of 1.5 $mm^{-1}$ based on measured curvature values in fig. S10, and that the composite structure of the three-segment bilayer would be formed by concatenating the shapes of each segment. The resulting design is shown in fig. S12.

**Simulation of Digit Gel Automata Curving.** Using data from bilayer DNA-co-polymerized gel characterization (fig. S10), we developed a numeric simulation to predict the final shapes of the gel strip. We characterized the radii of curvature (RoC) of bilayer gels and the changes in contour length resulting from different actuation combinations. A lookup into the resulting table of these values, indexed by each system type, set the radius of curvature and change in contour lengths of individual bilayers within a bilayer segment. To predict the shape of the contour of a strip consisting of multiple bilayer segments, we assumed that bilayer gels segments curved independently of one another. To simulate the curvature of gel automata, we began with a 1d array *segment_lengths* and a 2d array *identities* as inputs and generated a strip automata object. The *segment_lengths* array encoded the lengths of each segment, while *identities* encoded the types of systems in each segment. We then simulated the (up to) 16 possible states that a gel automaton could transform into given our four switchable DNA systems to produce 16 output images as follows. We first retrieved the values of the radii of curvature and change in contour from the tables referenced above to obtain a 1d array *rocs* for the radius of curvature of each segment and a 1d array *ctls* for the change in contour length of each segment. We then produced a final shape consisting of the concatenation of the final curves of each of the bent segments. These curves were produced by generating the next point along a composite curve using a numerical contour integration scheme. Specifically, starting at (x, y) = (0, 0) and q = 0, we used *delta* = 20 μm as a mesh size and an iterator object to generate the next (x, y) point as follows:

$$\Delta\theta = \frac{delta}{radius\ of\ curvature} \Longrightarrow \theta = \theta + \Delta\theta$$

$$\Delta x = delta \times sin\theta \Longrightarrow x = x + \Delta x$$

$$\Delta y = delta \times cos\theta \Longrightarrow y = y + \Delta y$$

The iterator generated new points within a given segment till the segment length limit $segment\ length \times (1 + \Delta contour\ length)$ for a segment was reached. The scheme then generated points for subsequent segments by continuing along the integration path. We next compiled 28 x 28-pixel images using the x, y points and applied a Gaussian filter (with sigma value = 0.8) to each generated image so that the images appeared similar in texture to the handwritten digits in the MNIST digits library (*37*). For more information and code demonstration, please visit the following GitHub repo: https://github.com/charliecharlie29/Deep-Learning-Guided-Genetic-Algorithm.git

**Convolutional Neural Network (CNN) for Autonomous Classifiers.** We compiled a convolutional neural network model with the TensorFlow library (Ver 2.4.3) and trained the model with a dataset containing the MNIST digit dataset and a dataset generated by strip automata geometry simulation described in "Simulation of Gel Digit Automata Curving". The generated dataset includes twenty-eight thousand human-labeled strip images. These images were either recognizable as one of the 0-9 digits (and thus were labeled with respective digits) or were considered "random squiggles", i.e., curves not representing any Arabic numeral, and were labeled as an eleventh category. Random squiggles were included as a class so that the model could determine whether a shape resembled any digits, in addition to quantifying the resemblance of a shape to the digit. We combined the generated dataset and the MNIST dataset into a combined dataset for training. The combined dataset consisted of ninety-eight thousand images and was split into train and test sets, which contained eighty-four thousand and fourteen thousand images, respectively. The data were normalized prior to training so that each pixel value lay between 0 and 1. The CNN model consisted of two convolutional layers with rectified linear unit (Relu) activation and max-pooling layers. Dropout layers were included to avoid overfitting, and a flattened layer was added prior to the fully connected layers. Two fully connected layers with Relu activation and a final output layer with softmax activation were present at the end of the network for classification. The model was compiled with categorical cross-entropy loss, trained using the Adam (Adaptive momentum estimation) optimizer with the default learning rate in Python's ScikitLearn. The trained network achieved ninety-eight percent accuracy on the test set by the end of training. Scripts used can be found at https://github.com/charliecharlie29/Gel-Automata-Directed-By-DNA-Codes

**Genetic Algorithm for Digit Gel Automata Parameter Search.** We developed a genetic algorithm to efficiently search through the large parameter space of bilayer strips to find designs for digit automata. The algorithm started with an initial population of automata designs generated from a random seed. Each design within the population was then simulated to find all possible geometric outputs of each of the 16 actuation combinations. The digit that each output resembled and its extent of resemblance are calculated using the network described in "Convolutional Neural Network (CNN) for Autonomous Classifiers". During the scoring process, all images were rotated at twenty different angles, and the image with the highest score as a digit was selected to represent the final class and the score of the image. The scores for each of the 16 actuation combinations were stored in a 2d array documenting what digits were formed and the score for each digit. A custom loss function is used to evaluate the fitness of each design:

$$loss = 5000 \times number\ of\ digits\ formed \times \sum_{i=0}^{9} \ln(1.001 - score\ for\ digit\ i)$$

The loss function computes the diversity and the similarity to real digits of the digits formed. Designs with output images that resemble a larger number of different, easily recognized digits are fitter according to this loss function and, therefore, more likely to be preserved in the population than those with fewer such output images. During the selection stage, 80% of the designs within the population are eliminated by selecting the designs with the 20% lowest (best) loss score to preserve. These preserved designs are sent into a mutation function to repopulate a new generation. The mutation was performed using the single-parent mutation method, where the genetic information of each descendant comes from a single survived design from the previous selection. During mutation, each design had a fifty percent chance to randomly update the strip segment lengths, preserving the activator pattern information. Otherwise, half of the regions in the activator pattern were mutated. Each survivor design generated four descendants, so the population returned to its original size after every round of selection and mutation. Finally, the algorithm iterated the cycle of population generation, selection, and mutation until reaching the generation limit, and output the optimized designs. For our even digit automata and odd digit automata search, we slightly tweaked the loss function and mutation function to obtain fabricable devices. We first included an additional rule within the mutation function to ensure new designs are within reasonable patterning steps to avoid generating designs that are overly complex and thus un-patternable. The number of fabrication steps was calculated as the sum of the number of unique activator systems in each layer. Patterns that require more than six fabrication steps were eliminated from consideration and either regenerated (if part of the initial population) or re-mutated from a parent (during subsequent rounds). We used this algorithm to search for an even digit automaton and an odd digit automaton, changing the loss functions for the two searches and deriving the final optimized outputs (fig. S14-15).

$$loss = 5000 \times number\ of\ digits\ formed \times \sum_{i=1,3,5,7,9} \ln(1.001 - score\ for\ digit\ i)$$

$$loss = 5000 \times number\ of\ digits\ formed \times \sum_{i=0,2,4,6,8} \ln(1.001 - score\ for\ digit\ i)$$

**Actuation of Letter and Digit Gel Automata.** As-made gel automata were transferred to a 1-inch diameter glass bottom petri dish for actuation and imaging. During monolayer and bilayer gel actuation, 1 ml of TAEM-Tween20 solution consisting of 60 μM DNA activators mix (99% growth activators, 1% growth terminators) was added to the petri dish. When further tuning of specific regions of the gel automata was needed, an additional 15 μM DNA activators mix was added to the solution as described in Fig. 4B. Between each actuation step, the old DNA solution was removed, TAEM-Tween20 buffer was added for 15 mins and removed before the DNA solution for the next step of actuation was added to the petri dish. Before each subsequent activation step, bright-field images were taken using a Hayear 4K Microscope Camera (HY-1070). For the letter gel automaton, the actuation process was recorded using the gel imager Syngene EF2 G: Box using the imaging protocol described above.

**Gel Automata Image Processing.** Images of gel automata actuation results were processed using MATLAB's Image Processing Toolbox 2022a. For each image, all major objects are recognized by brightness thresholding followed by a morphology closing, and the object near the center of the image was selected as the gel body. We then filtered individual images by dimming the background brightness while keeping the brightness of the gel body unchanged. Specifically, we created a binary mask that covered the gel body and dimmed the brightness of the region outside the mask to zero. We then matched the image histogram of all images to the first image (in the actuation process) histogram as a reference. For the letter gel automata actuation, the intensities of over-exposed regions were reduced by 15%.

**Supplementary Text S1: Processive depolymerization.**

**Designed mechanism of DNA crosslink polymerization.** The DNA crosslinks are in a double-stranded form with two toeholds (fig. S1a) inside the DNA gel automata structure. In the scheme for processive polymerization and depolymerization, growth activators are used in initiating gel growth. Each growth activator strand has one long toehold (domain a or domain c' in fig. 1a, pink or brown respectively in fig. S1a) and one short toehold (domain x' or y' in fig. S1a, emerald or green, respectively). Both the long and short toeholds are necessary to facilitate polymer chain growth. Initial growth proceeds identically to the forward-driven process described in (*17, 18*), with growth activators binding to the active sites of the polymer chain by two toeholds and undergoing four-way branch migration to extend the polymer chain. During polymerization, the single-stranded extensions on the growth activators (domains M and N in fig. S1a, yellow and magenta) are inert. These extensions are designed to serve as initiation sites for binding to the shrinking activators during shrinking.

**Designed mechanism of processive shrinking.** Processive shrinking activators are used with the goal of shrinking the hydrogels. The shrinking activators bind irreversibly to the M and N extensions on the growth activators that are incorporated into the polymerized chains (fig. S1c). Bound shrinking activators can reversibly branch migrate into the large toehold domains on the growth initiators (domains a:a' and c:c' in fig. S1a, pink and brown). When these large toeholds are bound to the shrinking activators, polymer chain shrinking becomes favored over polymer chain growth. The short toeholds (domains x:x' and y:y' in fig. S1a, emerald and green) at the active site of the polymer can then bind and initiate a four-way branch migration process initiated by a 3nt toehold to displace the most recently bound growth activators (fig. S1c). This process can repeat until processive reactions have displaced all bound growth activators from the polymer. There is also a side depolymerization reaction initiated with a 0nt toehold (fig. S1d), which we assume would be much slower in rate.

When we conducted a processive shrinking experiment (fig. S2); we did not observe any gel shrinking when the processive shrinking activators were added. We hypothesized that this null result might be due to the slow kinetics of the reactions.

The following argument supports this hypothesis. Assuming the reactions to happen with a 100μM effective concentration of activators and using the rate from (*32*), then

$$Processive\ depolymerization\ rate = 10^{-1}\mathrm{M}^{-1}\mathrm{sec}^{-1} \times 100\mu\mathrm{M} = 10^{-5}\mathrm{sec}^{-1}$$

$$Processive\ depolymerization\ time\ constant = \frac{1}{Processive\ depolymerization\ rate} \approx 30\ \mathrm{hr}$$

For the side depolymerization reaction initiated with a 0nt toehold, assuming the reactions happen with the same effective concentration and using the rate from (*32*):

$$Side\ depolymerization\ reaction\ rate = 10^{-2}\mathrm{M}^{-1}\mathrm{sec}^{-1} \times 100\mu\mathrm{M} = 10^{-6}\mathrm{sec}^{-1}$$

$$\mathit{Side\ depolymerization\ reaction\ time\ constant} = \frac{1}{\mathit{Processive\ depolymerization\ rate}} \approx 300\ \mathrm{hr}$$

Our estimated calculations, based on rate constants drawn from (*32*), suggest that it takes approximately 30 hours to displace each growth monomer from the polymer chain in a processive manner. This estimate indicates that it would take weeks to depolymerize the entire polymer chain, which is well beyond the timeframe of our experimental observations and thus would explain why we were unable to observe shrinking for this design. Altering the design to improve the rate was not straightforward. The reaction rate constants for toehold-mediated four-way branch migration process are generally smaller than the reaction rate constants for three-way branch migration rates. The reaction rate constants for toehold-mediated four-way branch migration process are also greatly dependent on the sequence designs, and do not increase significantly by simply making the toeholds longer (*32*). Changing the sequence lengths of x and y domains will not affect the activation energy required to break the bonds of polymerized crosslinks, but only the final energy state, and thus the effect on reaction rate is limited. Additionally, the processive shrinking design is also affected by the application of strain across the polymer crosslink applied by the parent gel matrix, which is not accounted for by the model in (*32*).

**Redesign for fast, parallel breakage of polymerized crosslinks.** We thus chose to redesign the sequences of the shrinking activators to reach into the branch migration domains of the growth activator hairpins (domain b in fig. S1a, dark blue; design schematics in Fig. 1A and fig. S1e). After this three-way branch migration step occurs, all the growth activators are fully disconnected from each other and from the root DNA crosslinks, which depolymerizes the DNA crosslinks quickly.

**Supplementary Text S2: Multi-step DNA gel automata photopatterning process.**

During photolithography for hydrogel fabrication (*38*), a lithography chamber is first assembled to hold the liquid prepolymer. The chamber consists of a photomask, a substrate, and two spacers with a thickness determining the hydrogel dimension. To create a multi-step hydrogel photopatterning process that could be used reliably over up to five steps, we substituted an identical photomask for the glass substrate so that after the first step of photopatterning, the hydrogel remained on a substrate with alignment markers (fig. S11). The alignment markers are crucial in the following patterning steps as the alignment accuracy plays an essential role in the final morphology. Misalignment between layers or gel regions can result in bending or curling of the hydrogel strip after hydration and create errors in the shape of gel automata actuation. Also, undesired deformation of the hydrogel strip can occur during hydration when the hydrogel strip has thickness variations, especially when the aspect ratio is large. Although we used the same spacers to control the height of each hydrogel region during each hydrogel patterning step, the existing hydrogel layer can be hydrated by the pre-gel solution, causing it to swell up and take more space, which results in a difference in thickness between the first and second layers. Hence, we included a rinsing and gently blow-drying process between each patterning step. We also sought to keep the alignment process fast to limit this hydration effect. Even though the photomasks used are made from 1.5 mm-thick glass slides and are, therefore, quite rigid, there is still slight bending of slides when they are clamped on two sides and sandwich the spacers during UV exposure. This bending will affect the height homogeneity across the chamber in different positions and cause thickness variations in the patterned structures. We therefore optimized the photomask design to utilize the central space of the photomasks, where the effects of bending are the most minor, to manufacture structures with consistent heights. We reduced material loss due to undesired gel lift-off on Cr masks by coating CYTOP (a hydrophobic fluoropolymer) onto the substrate to which high MW PEG tends to adhere and by cleaning the photomask with DI-water right before use. All multi-step photopatterning masks were designed to present features 1.5 times larger in length than those that were designed by computer programs to compensate for the potential curvature reduction in subsequent actuation cycles and the constraining effect from adjacent gel segments. We also varied the sizes of the overlapping areas of neighboring hydrogel regions to be between 100 and 200 µm. These sizes were chosen to balance the amounts of material used for patterning, the robustness of the junction of devices being patterned, and the ensuing difficulty of aligning masks during different patterning steps.

**Supplementary Text S3: Programmable Imaging System (Pi-Imager) for Time Lapse Fluorescence Image Capture.**

To acquire time-lapse fluorescence images for our hydrogel, we designed a programmable imaging system using Raspberry Pi, an Arducam camera, and optical components.

The imaging systems, termed “Pi-Imager,” consisted of the following:

1. Raspberry Pi computer for autonomous control
2. Arducam Pi camera for image capturing
3. Thorlabs LED array lamps, filter sets, and motor for multiple fluorescence imaging
4. Thorlabs optical enclosure and caging system for structure holding

The imaging system is connected and controlled through a Raspberry Pi Linux computer to control an LED array lamp for fluorescence excitation and a filter motor that switches between different corresponding fluorescence spectra for excitation. Images are then captured by the connected Arducam Pi camera and saved to the Raspberry Pi computer for further image analysis. Developed scripts for system control can be found here: https://github.com/charliecharlie29/Hydrogel-Automata-Directed-By-DNA-Codes.

Instrument specifics:

1. Raspberry Pi 4 4GB Model B with 1.5GHz 64-bit quad-core ARMv8 CPU (4GB RAM)
2. Arducam for Raspberry Pi Camera, Interchangeable CS Mount Lens for Pi 4, 3, 3B+, 5MP OV5647 1080P
3. HAYEAR Monocular Max 180x Zoom C-Mount Glass Lens Adapter F/Industry Microscope Camera Objective
4. Youngneer 5v Relay Board Raspberry Arduino Relay Module 1 Channel Opto-Isolated High or Low Level Trigger
5. MCIGICM 50pcs ic dip lm358 Operational Amplifier Dual op-amp (Pack of 50pcs)
6. ALITOVE 5V 60A 300W Power Supply Transformer Adapter Converter AC110V/220V to DC 5V 60amp LED Driver for WS2812B WS2811 WS2801 APA102 LED Strip Pixel Light CCTV Camera Security System
7. Thorlabs Multi-Position Sliders, ELL9, ED1-C50-MD, SM1L03
8. Thorlabs Filter Sets, MDF-TOM, MDF-YFP
9. Thorlabs LED Array Light Sources, LIUCWHA, LIU-PS, CAB-LEDD1, CON8ML-4
10. Thorlabs Black Hardboard Enclosures, XE25C9, XE25C9D, MB1824

11. Thorlabs Cage System, ER6, CP33, TR6, RA90

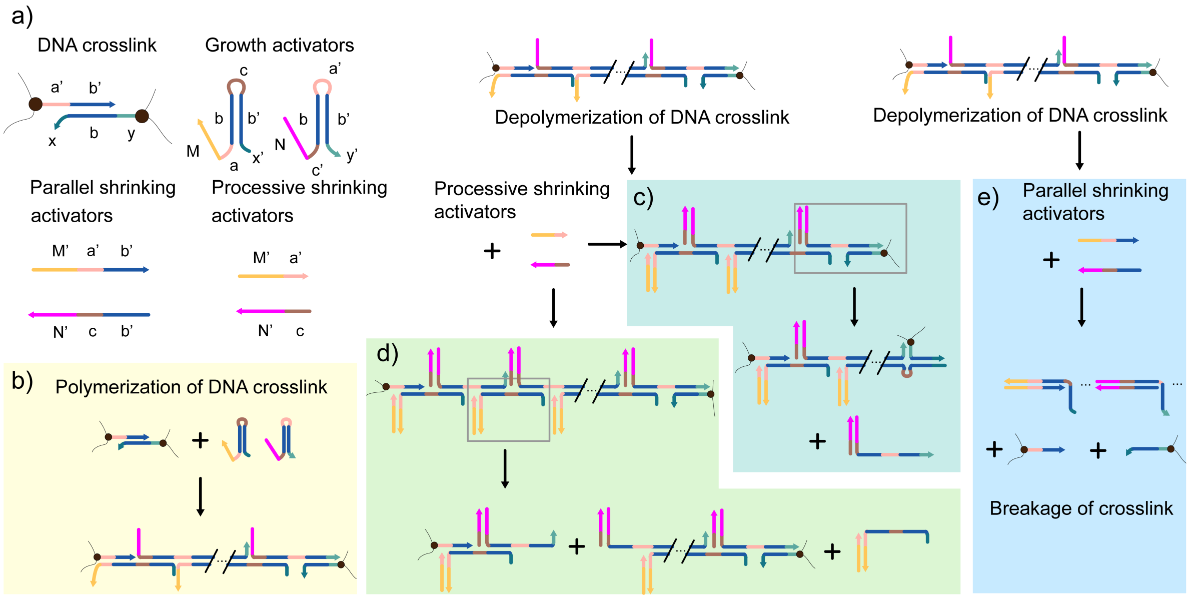


**Figure S1. Design of a processive shrinking directive and a parallel shrinking directive for DNA hydrogel shape change and their mechanisms.** A detailed discussion of the design can be found in Supplementary Text S1. DNA sequences used are listed in Table S1. Processive Shrinking Directive strands use the acronym PSD. a) DNA strands and complexes. Domain colors: a:a' pink; b:b' dark blue; c:c' brown; x:x' emerald; y:y' green; M:M' yellow; N:N' magenta. b) Polymerization of DNA crosslink. c, d) Processive depolymerization. e) Parallel depolymerization.

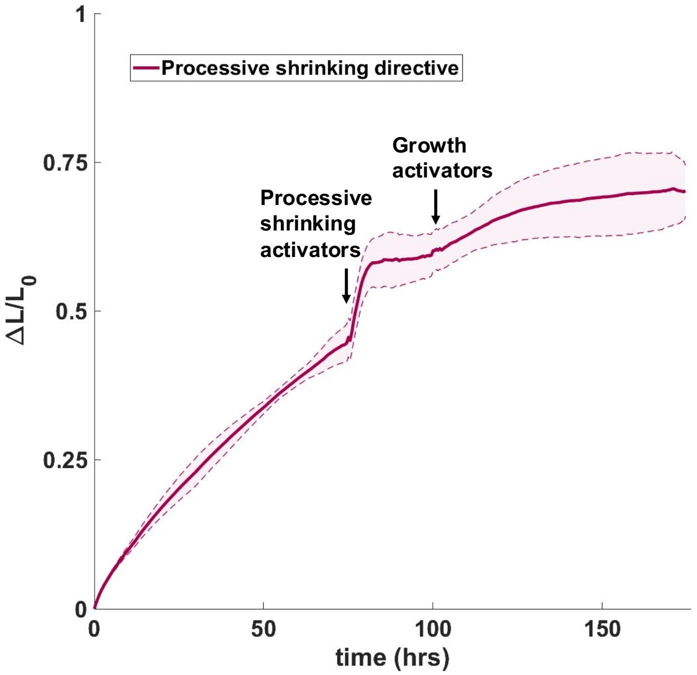


**Figure S2. Actuation using processive shrinking directives.** Initially, PEG-co-DNA hydrogels were each added to 150 µL of TAEM-Tween20 solution containing 20 µM growth activators (Processive Shrinking Directive (PSD) strands, sequences listed in Table S1). After 75 hours of growth, the activator DNA solution was removed, then 100 µL TAEM-Tween20 was added for 15 mins and removed, and 150 µL of 20 µM processive shrinking activator DNA sequences mixed with TAEM-Tween20 was added. After 25 hours, the activator DNA solution was removed, then 100 µL of TAEM-Tween20 was added for 15 mins and removed, and 150 µL TAEM-Tween20 solution containing 20 µM growth activators was added to the gel, as in the first step.

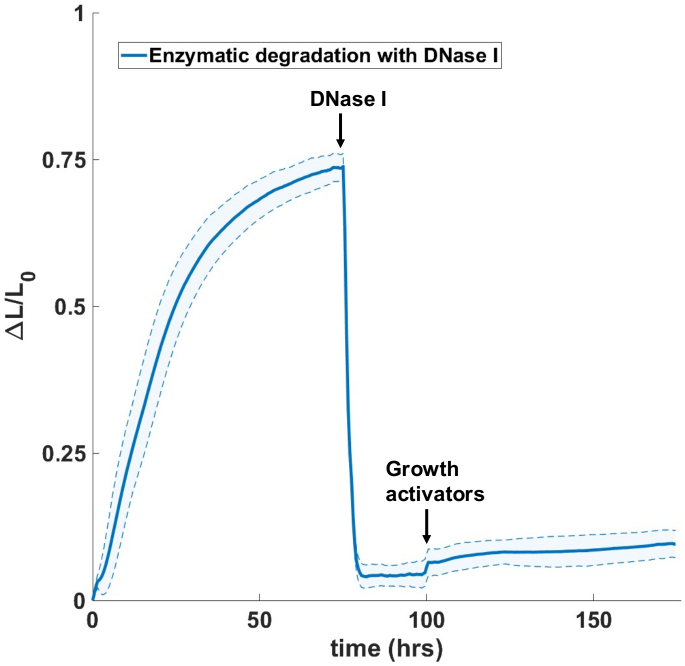


**Figure S3. Enzymatic degradation shrinks DNA hydrogels; this shrinking is not reversible.** Initially, PEG-co-DNA hydrogels were each added to 150 μL of TAEM-Tween20 solution containing 20 μM growth activators (Enzymatic degradation strands, sequences listed in Table S1). After 75 hours of growth, the DNA solution was removed, and 100 μL TAEM-Tween20 was added for 15 mins and removed. Then the gels were treated with two units of RNase-free DNase I (New England BioLabs, M0303) in 100 μL DNase I reaction buffer at room temperature. After 25 hours, the hydrogels were rinsed with TAEM-Tween20 three times and added to 100 μL of TAEM-Tween20 for 24 hours, then treated with the same TAEM-Tween20 solution containing 20 μM growth activators as in the first step.

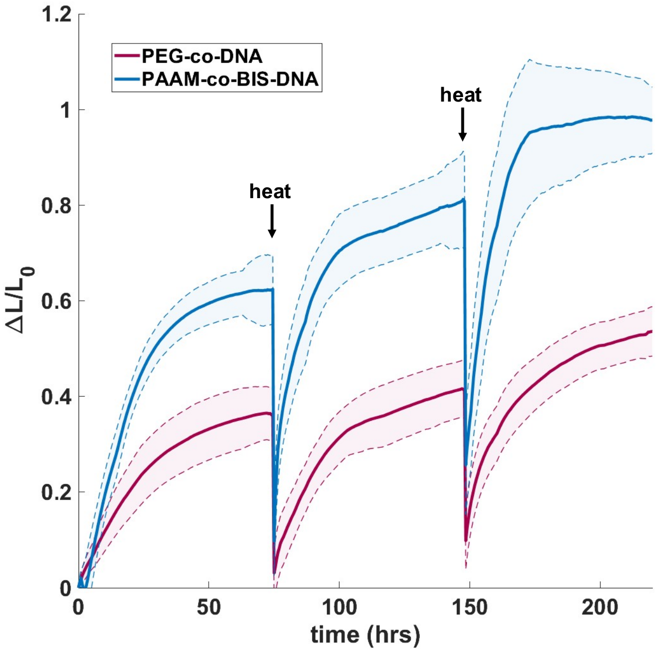


**Figure S4. Changes in the size of PEG-co-DNA and PAAM-co-BIS-DNA hydrogels during cycles of growth at room temperature followed by heating to 95ºC.** Initially, both types of hydrogels were added to 150 µL of TAEM-Tween20 solution containing 20 µM growth activators (Heat-induced reversibility strands, sequences listed in Table S1). After 72 hours of growth, the DNA solution was removed, and 100 µL TAEM-Tween20 was added to the gels for 15 mins at 95°C in a PCR machine and then cooled down to room temperature. We later removed the TAEM-Tween20 and treated the gels with 150 µL of fresh 20 µM growth activators with TAEM-Tween20 again. The same heat treatment happened one more time after 72 hours. The gels were monitored under fluorescence microscopy except during heating.

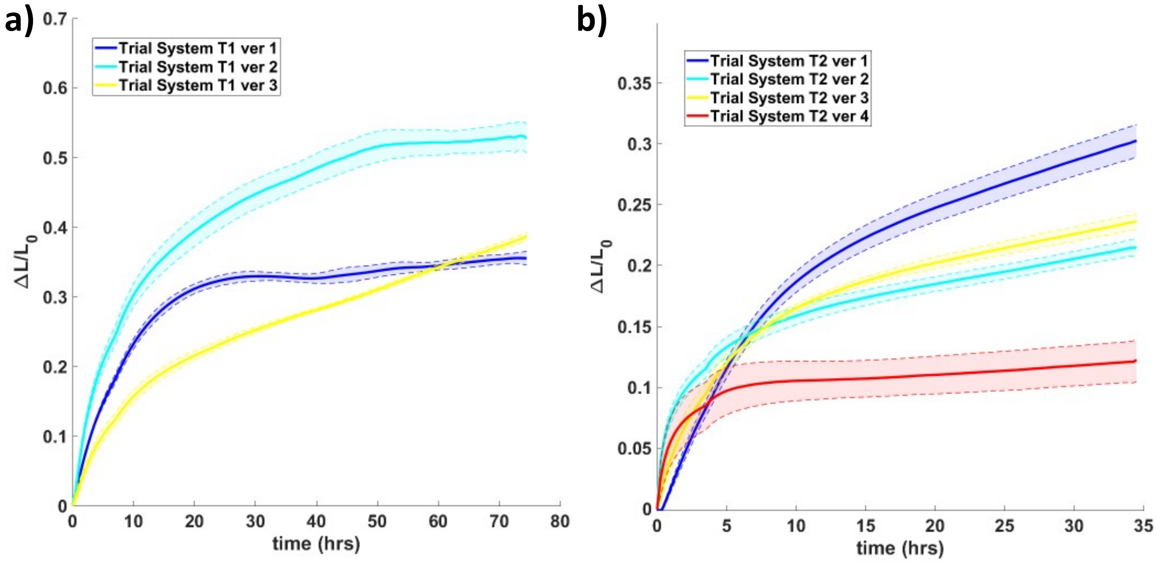


**Figure S5: Sequence dependence of growth activators.** Three and four versions of trial systems, a) T1 and b) T2, were designed based on sequences from (*2*) (Trial System T1 and T2, sequences listed in Table S1). For each trial system, we designed the growth activators with different M and N domain sequences, as shown in fig.S1a. PEG-co-DNA(MW20k) hydrogels were each added to 150 μL of TAEM-Tween20 solution containing 20 μM growth activators. The sequence difference between each version of the growth activators was 15 bases/71 bases = 21.12%. The results suggest the differences in sequence significantly altered the growth speed and final extent of growth.

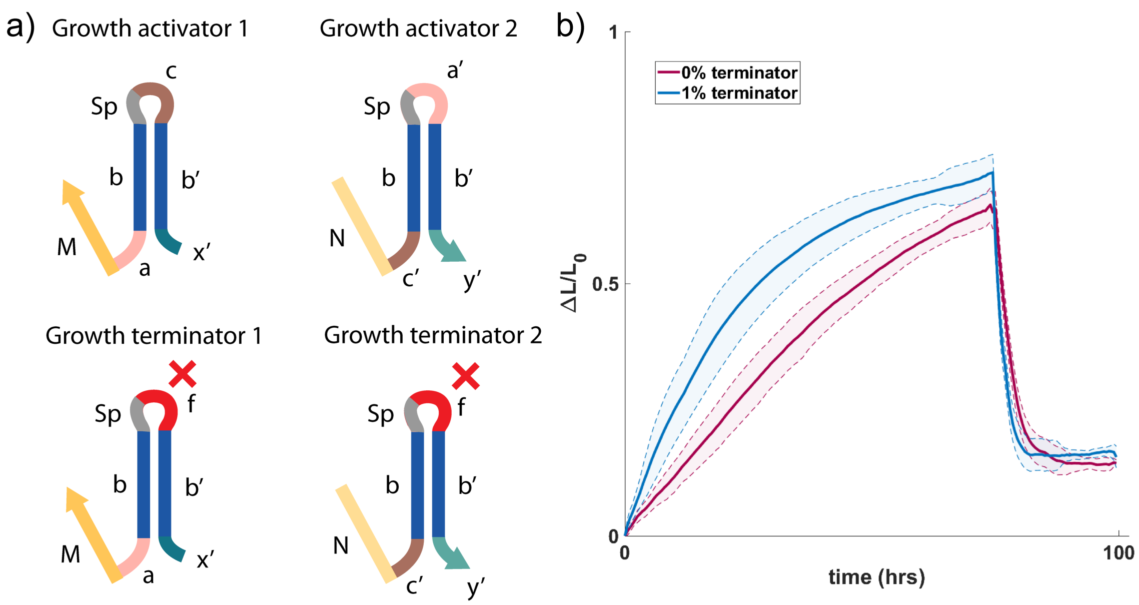


**Figure S6. Kinetics of gel growth and shrinking when growth activators are used with or without growth terminators.** a) Schematics of different sequence designs. b) PEG-co-DNA hydrogels were first treated with 150 μL of 60 μM growth activators containing 0% and 1% growth terminator in TAEM-Tween20, respectively. After 72 hours of growth, the DNA solution was removed, and 100 μL TAEM-Tween20 was added for 15 mins then removed. The same volume and concentration of shrinking activator strands were then added to the hydrogels. Sequences used are listed in Table S1 as System I strands.

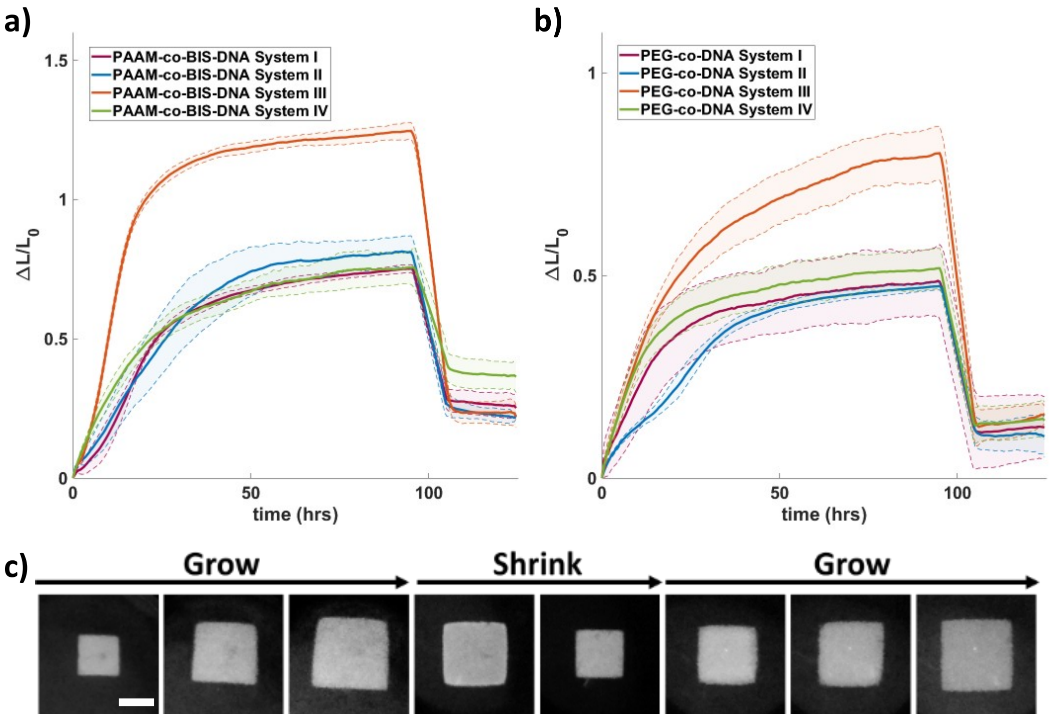


**Figure S7: DNA-driven growth and shrinking of PAAM-co-BIS-DNA and PEG-co-DNA gels crosslinked by different sequences in response to their respective activators.** a) Kinetics of shape change of PAAM-co-BIS-DNA hydrogels in response to System I, II, III, and IV DNA activators. b) Kinetics of growth and shrinking of PEG-co-DNA hydrogel in response to System I, II, III, and IV DNA activators. c) Time-lapse fluorescence micrographs of a PEG-co-DNA System III hydrogel after adding growth activators at 0 hr, 25 hr, 100 hr; after adding shrinking activators at 0 hr, 12 hr; and after adding growth activators again at 0 hr, 25 hr, 100 hr. Scale bar, 1 mm. Sequences used are listed in Table S1 as System I, II, III, and IV strands, [DNA activators] = 60 μM in TAEM-Tween20 with 1% growth terminators. The solution exchange process is as previously described in Methods.

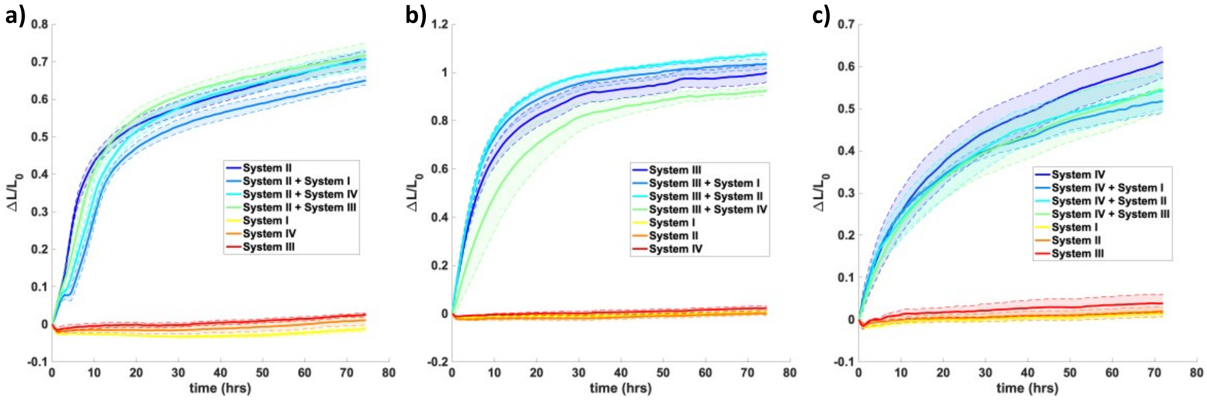


**Figure S8. There is minimal crosstalk between different systems of growth activators and hydrogels with certain DNA crosslinks.** a) System II, b) System III, and c) System IV PEG-co-DNA(MW20k) hydrogels were respectively treated with specific sets of DNA growth activators mix in TAEM-Tween20 as shown in the figure legends (150 µL, 60 µM). DNA sequences are listed in Table S1 as System I, II, III, and IV strands.

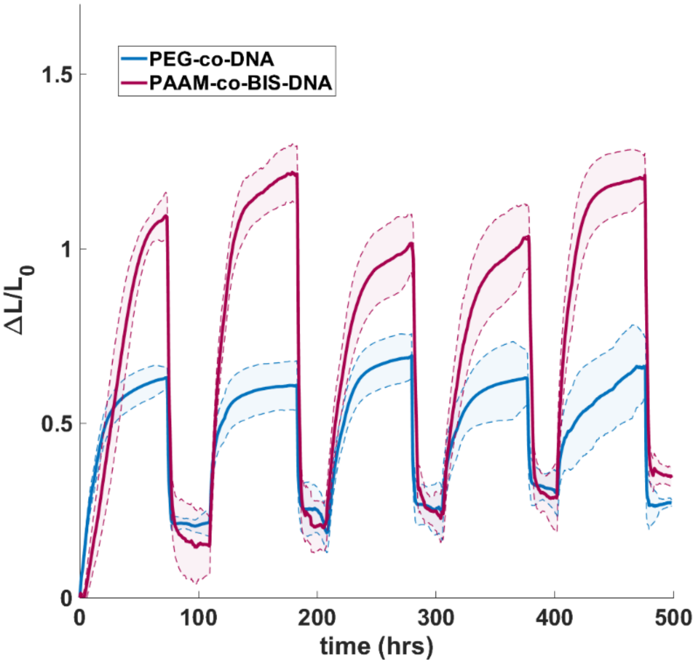


**Figure S9. Size change in PEG-co-DNA(MW20k) and PAAM-co-BIS-DNA hydrogel during five cycles of growth and shrinking.** Each actuation cycle consisted of 1) adding 150 µL growth activators to a final concentration of 60 µM with TAEM-Tween20 buffer for 72 hours; 2) removing the DNA solution, adding 100 µL TAEM-Tween20 for 15 mins, and then removing the solution; 3) adding 150 µL shrinking activator strands to a final concentration of 60 µM for 48 hours. 4) removing the DNA solution, adding 100 µL TAEM-Tween20 for 15 mins, and then removing the solution. DNA sequences used are listed in Table S1 as System II_v1 strands.

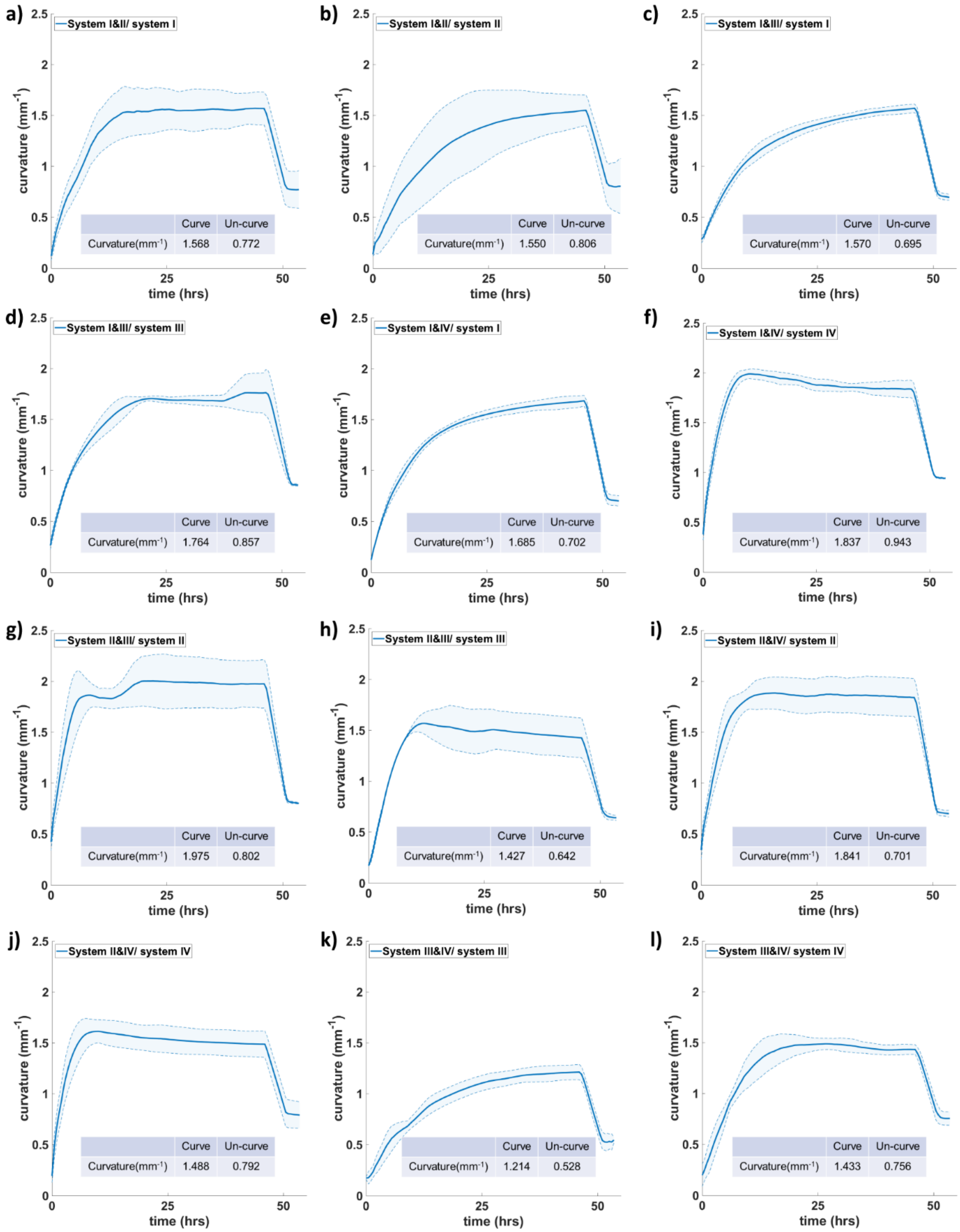


**Figure S10. Measurement of curvatures of gel bilayers with different DNA system combinations.** Curvature changes of PEG-co-DNA bilayer hydrogel treated with 'curve' and 'uncurve' instruction for every system combination shown in the figure legend. Curvature values are at steady states.

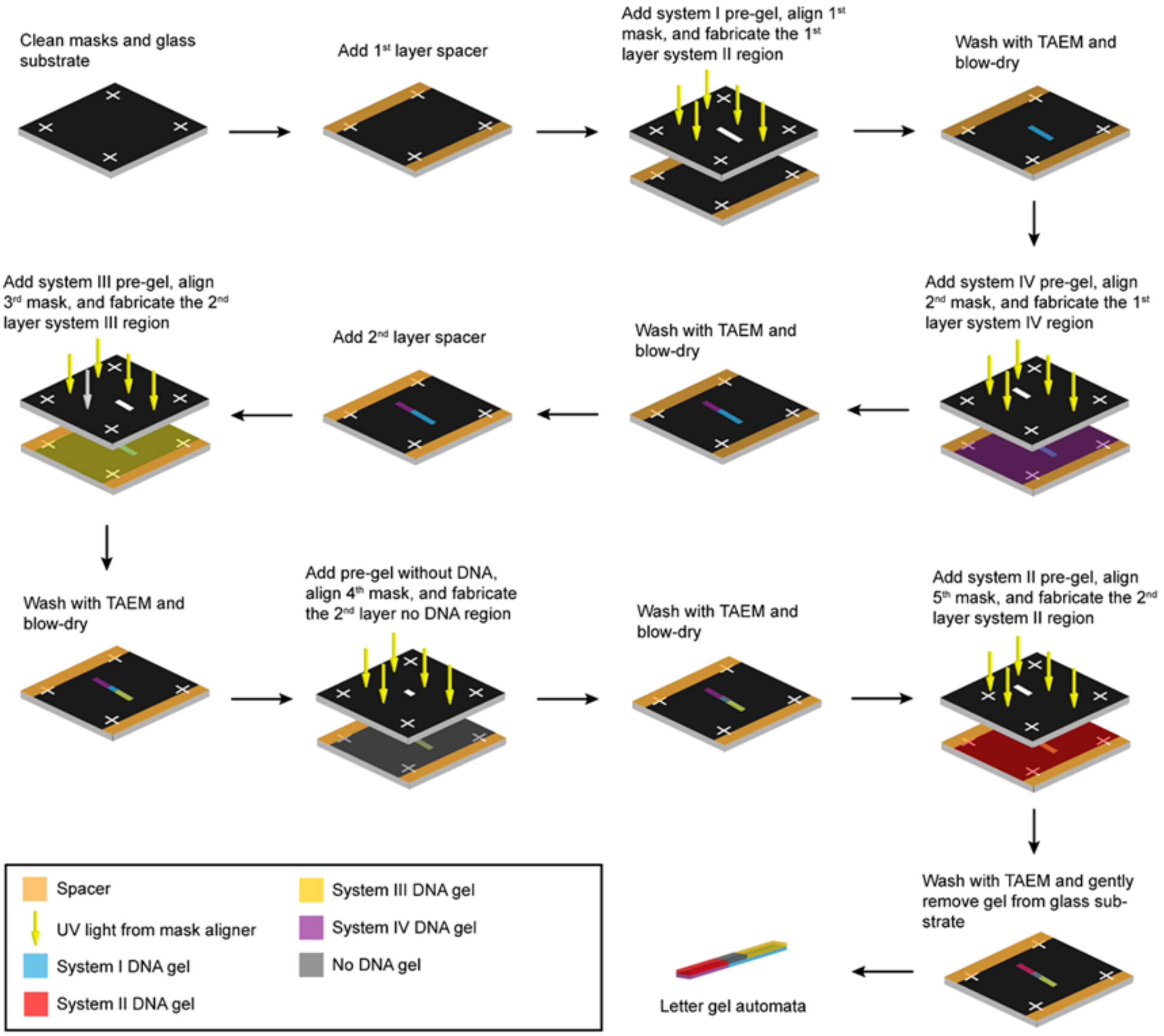


**Figure S11. Workflow for letter gel automata fabrication.** Parameters and detailed descriptions of the fabrication process can be found in the Methods and Supplementary Text S2.

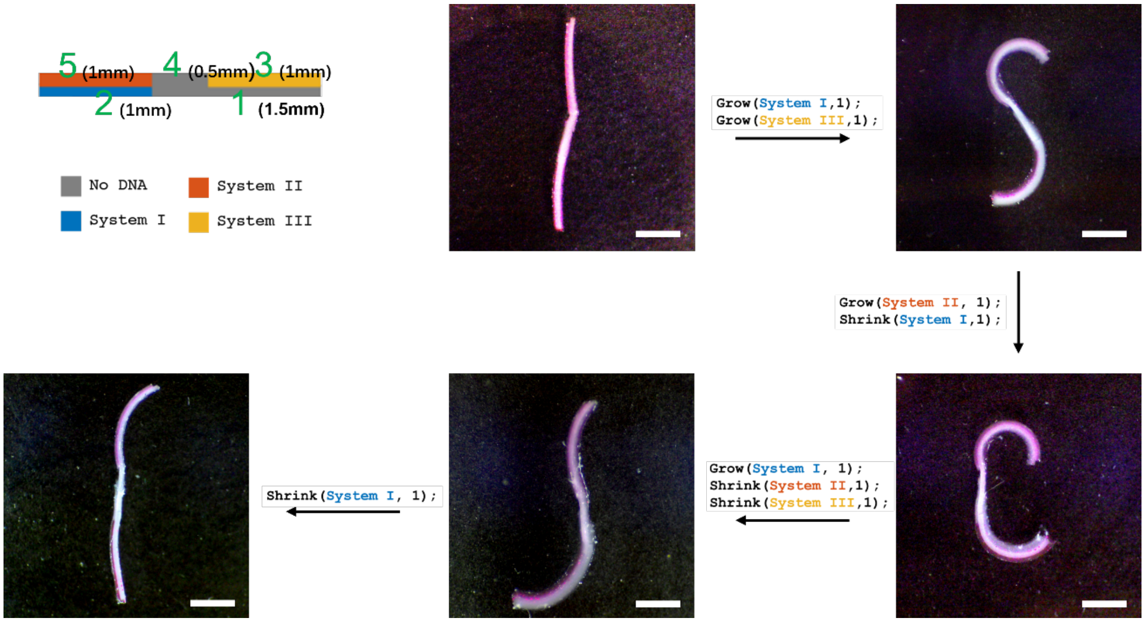


**Figure S12. Version 1.0 Letter gel automata.** Bright-field images of a V1.0 letter automaton before the start of actuation and after each step of the actuation program. The designed dimensions of each region of the gel automata from 1 to 5 are 1.5 mm, 1 mm, 1 mm, 0.5 mm, and 1 mm, respectively. Scale bars, 2 mm.

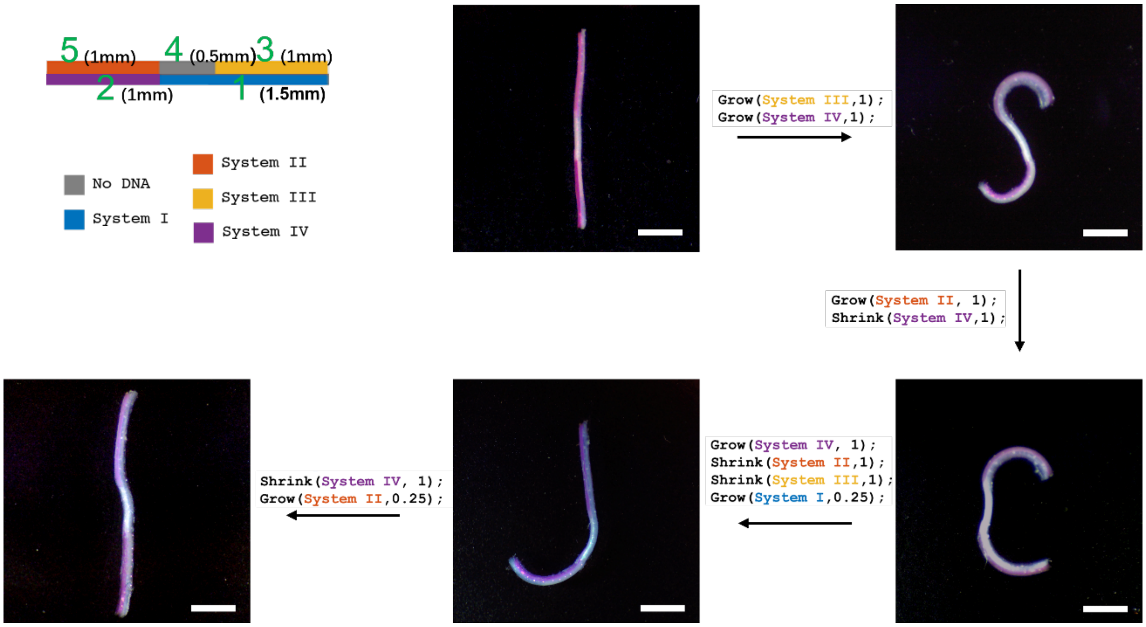


**Figure S13. Version 1.1 Letter gel automata.** V1.1 design and actuation program for letter gel automata with bright field images of letter gel after each program step. The designed dimensions of each region of the gel automata from 1 to 5 are 1.5 mm, 1 mm, 1 mm, 0.5 mm, and 1 mm, respectively. Scale bars, 2 mm.

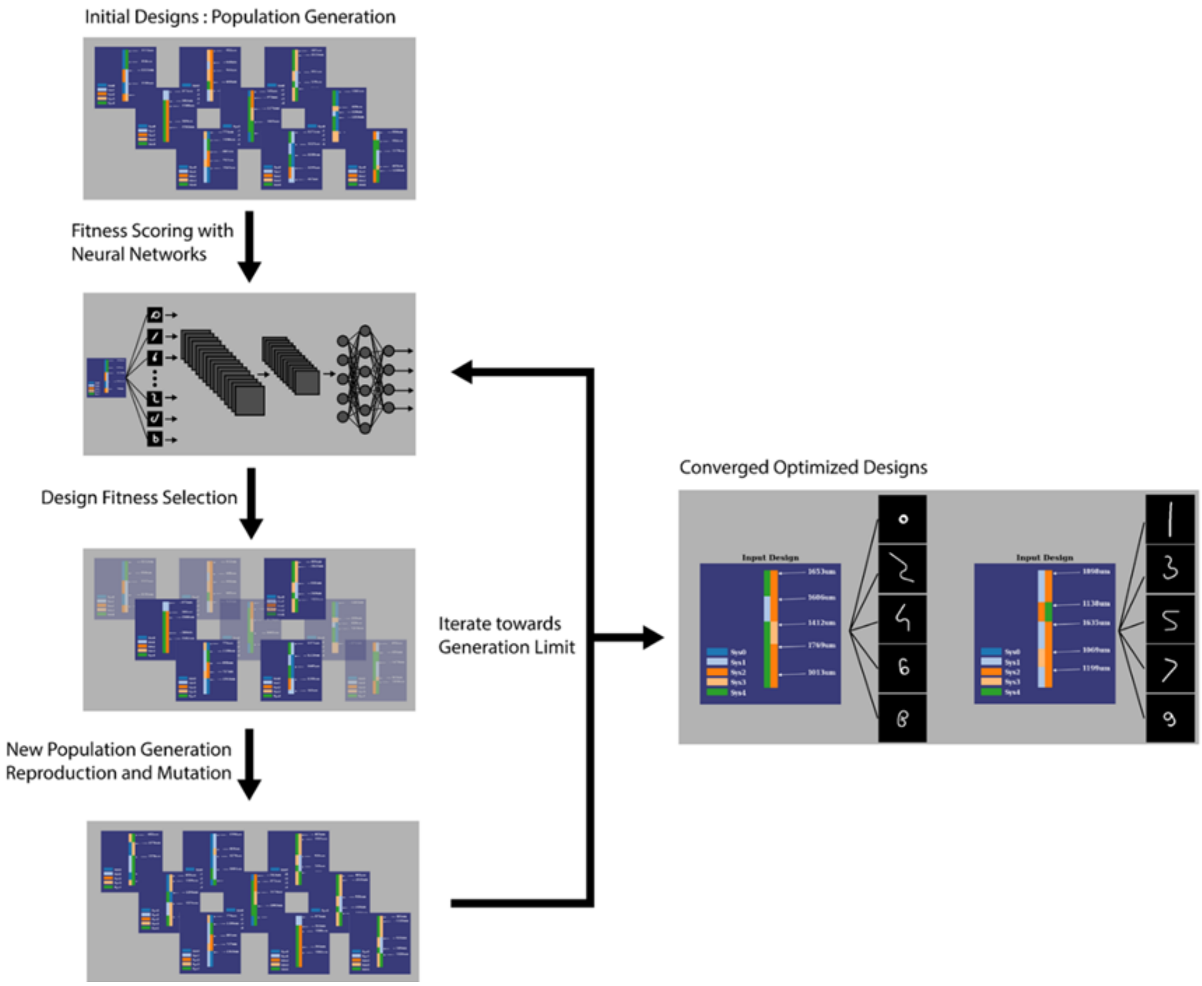


**Figure S14. Digit gel automata design process.** Detailed descriptions of the correlational neural network and design process can be found in the Methods section. The code used is available at: https://github.com/charliecharlie29/Hydrogel-Automata-Directed-By-DNA-Codes.

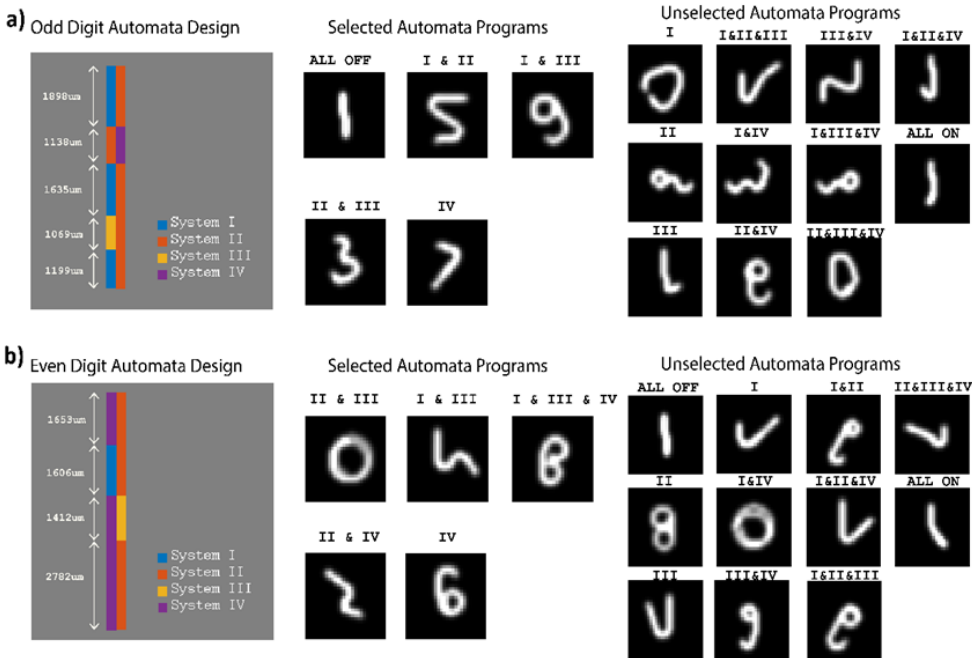


**Figure S15. Design and output simulations of a) odd digit and b) even digit gel automata.** The digits that the automata were designed to form from selected automata programs shown in the figure all received scores of more than 0.999 (max 1) from the neural network classifier.

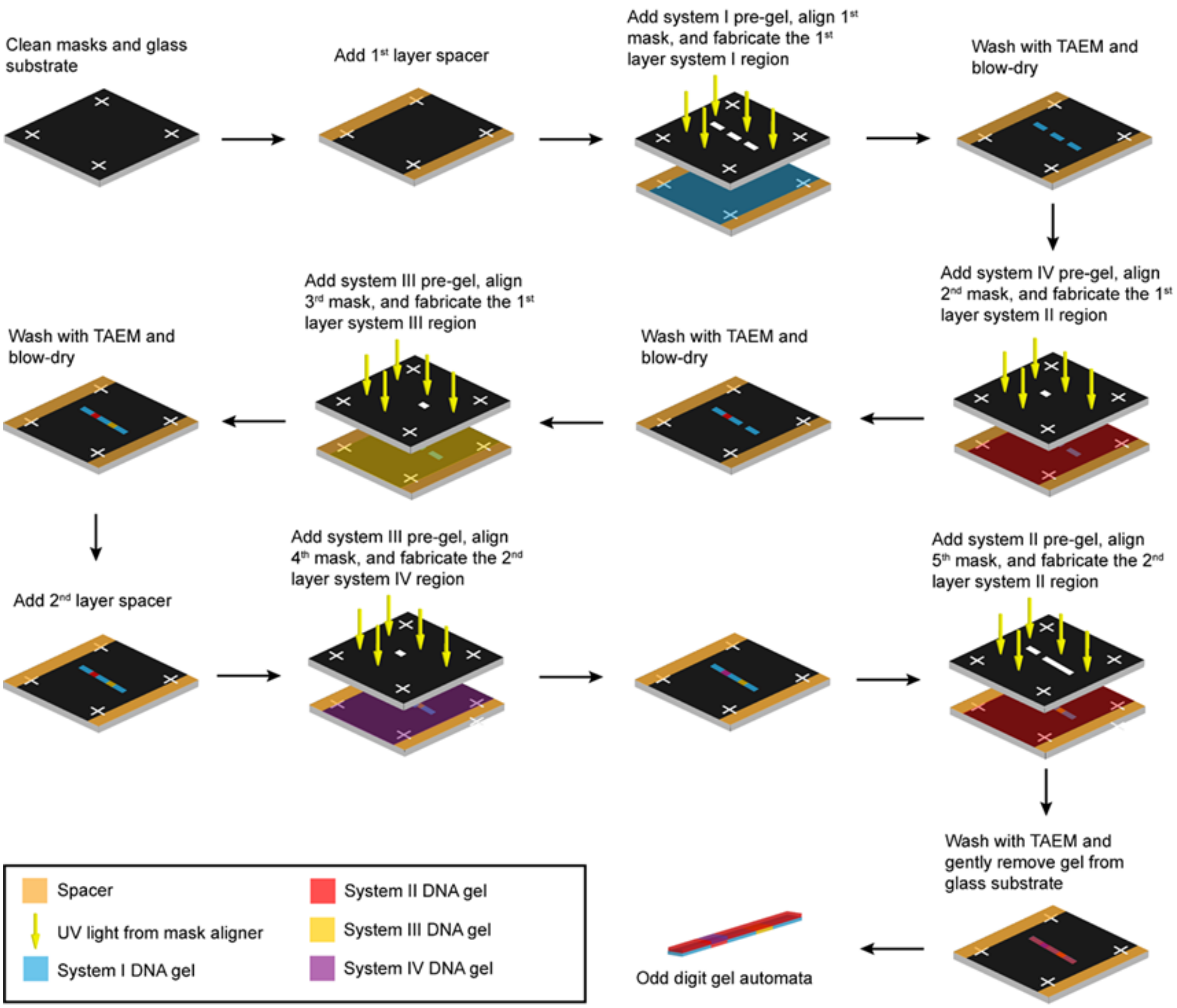


**Figure S16. Odd digit gel automata fabrication process.** Parameters and detailed descriptions of the fabrication process can be found in the Methods and Supplementary Text S2.

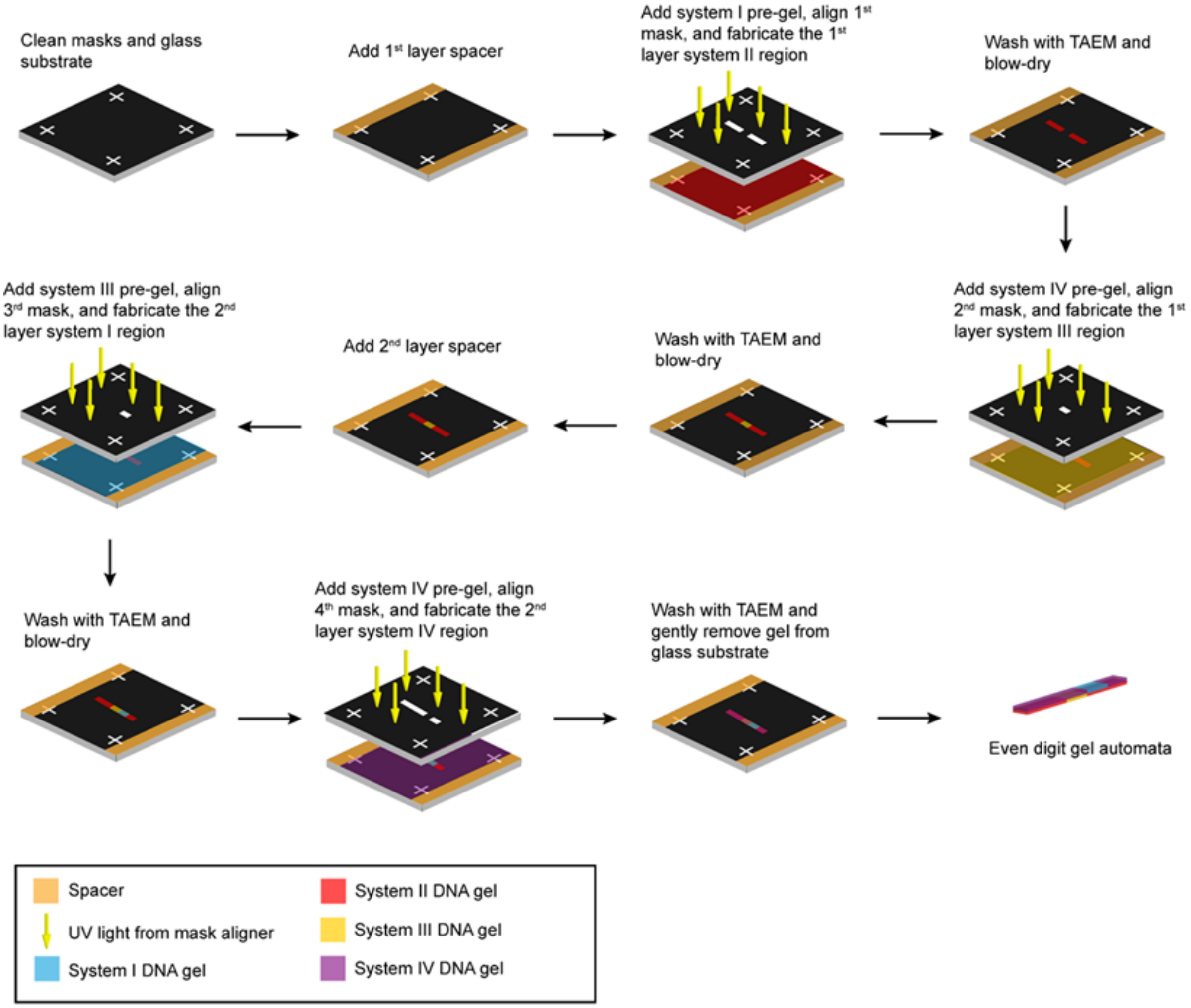


**Figure S17: Even digit gel automata fabrication process.** Parameters and detailed descriptions of the fabrication process can be found in the Methods and Supplementary Text S2.

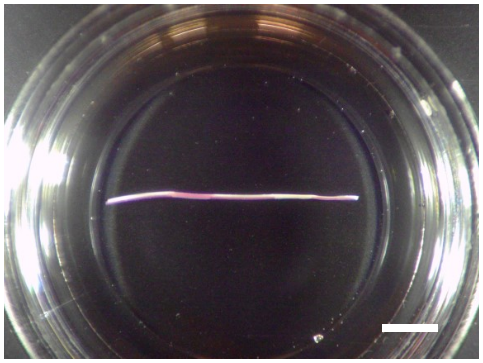

**Figure S18. Bright-field image of an as-made odd digit gel automata.** The resulting structure has an initially straight side view, as assumed in the design process. Scale bar, 5 mm.

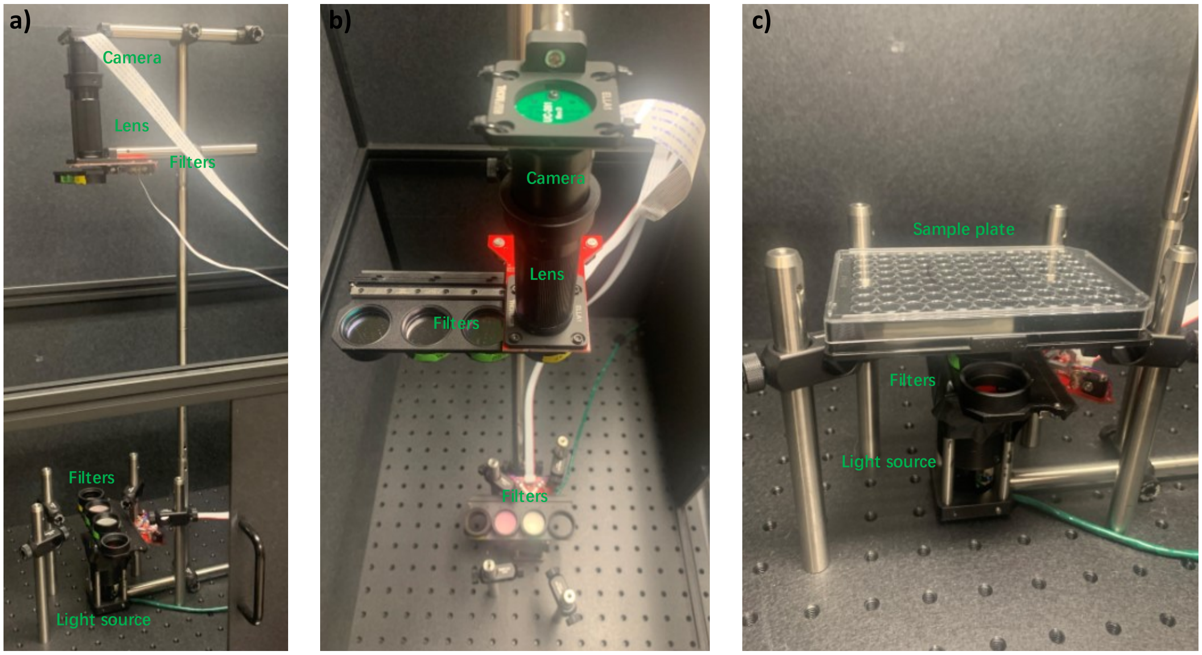


**Figure S19. Programmable Imaging System (Pi-Imager) for Time Lapse Fluorescence Images Capture.** We designed a programmable imaging system using Raspberry Pi, Arducam camera, and optical instruments acquired from Thorlabs to build an automated fluorescent imaging system tailored for our experimental needs. The Pi-Imager setup and scripts used can be found in Supplementary Text S3.

**Table S1. DNA sequences.** All strands were ordered from IDT in their lyophilized form, resuspended with TAEM solution, and stored at -20° C until use. /5ACryd/ indicates an Acrydite modification.

| Name | Sequence |
|---|---|
| System I | |
| System I_A | /5ACryd/CCTAAGTTCGCTGTGGCACCTGCACG |
| System I_R | /5ACryd/CAACGTGCAGGTGCCACAGCGTGG |
| System I_growth activator 1 | CCACGCTGTGGCACCTGCACGCACCCACAGCCATCGTGCAGGTGCCACAGCGAACTTAGGATGATTGTGTATAGT |
| System I_growth terminator 1 | CCACGCTGTGGCACCTGCACGTAAACATAGCCATCGTGCAGGTGCCACAGCGAACTTAGGATGATTGTGTATAGT |
| System I_growth activator 2 | AGTTAAGAGAATGATTGTGGGTGCGTGCAGGTGCCACAGCGGCCATCCTAAGTTCGCTGTGGCACCTGCACGTTG |
| System I_growth terminator 2 | AGTTAAGAGAATGATTGTGGGTGCGTGCAGGTGCCACAGCGGCCATTATTACTACGCTGTGGCACCTGCACGTTG |
| System I_shrinking activator 1 | ACTATACACAATCATCCTAAGTTCGCTGTGGCACCTGCACG |
| System I_shrinking activator 2 | CGCTGTGGCACCTGCACGCACCCACAATCATTCTCTTAACT |
| System II | |
| System II_A | /5ACryd/CTCTGTCTGCCTACCACTCCGTTGCG |
| System II_R | /5ACryd/ATTCGCAACGGAGTGGTAGGCTTT |
| System II_growth activator 1 | AAAGCCTACCACTCCGTTGCGGAACCTCCCTAAACGCAACGGAGTGGTAGGCAGACAGAGGTAAGGTAAGATAGG |

| | |
|---|---|
| System II_growth terminator 1 | AAAGCCTACCACTCCGTTGCGTCAAGCCCCTAAACGCAACGGAGTGGTAGGCAGACAGAGGTAAGGTAAGATAGG |
| System II_growth activator 2 | GGGTAGTGTGATGTGGGAGGTTCCGCAACGGAGTGGTAGGCCTAAACTCTGTCTGCCTACCACTCCGTTGCGAAT |
| System II_growth terminator 2 | GGGTAGTGTGATGTGGGAGGTTCCGCAACGGAGTGGTAGGCCTAAAAATCGTCCGCCTACCACTCCGTTGCGAAT |
| System II_shrinking activator 1 | CCTATCTTACCTTACCTCTGTCTGCCTACCACTCCGTTGCG |
| System II_shrinking activator 2 | GCCTACCACTCCGTTGCGGAACCTCCCACATCACACTACCC |
| System III | |
| System III_A | /5ACryd/CTATCTATCCATCACCCTCACCTTAC |
| System III_R | /5ACryd/GGTGTAAGGTGAGGGTGATGGTAA |
| System III_growth activator 1 | TTACCATCACCCTCACCTTACTTGTAGATTTTTTGTAAGGTGAGGGTGATGGATAGATAGGGTAGGTGAATGGGA |
| System III_growth terminator 1 | TTACCATCACCCTCACCTTACCTCTCCACTTTTTGTAAGGTGAGGGTGATGGATAGATAGGGTAGGTGAATGGGA |
| System III_growth activator 2 | TATGAGTGAGTTAGGATCTACAAGTAAGGTGAGGGTGATGGTTTTTCTATCTATCCATCACCCTCACCTTACACC |
| System III_growth terminator 2 | TATGAGTGAGTTAGGATCTACAAGTAAGGTGAGGGTGATGGTTTTTACGAGCCTCCATCACCCTCACCTTACACC |
| System III_shrinking activator 1 | TCCCATTCACCTACCATAGATAGCCATCACCCTCACCTTAC |

| | |
|---|---|
| System III_shrinking activator 2 | CCATCACCCTCACCTTACTTGTAGATCCTAACTCACTCATA |
| System IV | |
| System IV_A | /5ACryd/CTACCACTCCACTCACACTCCACTCC |
| System IV_R | /5ACryd/GGTGGAGTGGAGTGTGAGTGGGAT |
| System IV_growth activator 1 | ATCCCACTCACACTCCACTCCCGCTCGCCTAATAGGAGTGGAGTGT GAGTGGAGTGGTAGGTTTAGGTGAGGTGG |
| System IV_growth terminator 1 | ATCCCACTCACACTCCACTCCGTGCTGGTTAATAGGAGTGGAGTGT GAGTGGAGTGGTAGGTTTAGGTGAGGTGG |
| System IV_growth activator 2 | GTTGTAAGTGAGAGTGGCGAGCGGGAGTGGAGTGTGAGTGGTAAT ACTACCACTCCACTCACACTCCACTCCACC |
| System IV_growth terminator 2 | GTTGTAAGTGAGAGTGGCGAGCGGGAGTGGAGTGTGAGTGGTAAT AAAGGCGTCCCACTCACACTCCACTCCACC |
| System IV_shrinking activator 1 | CCACCTCACCTAAACCTACCACTCCACTCACACTCCACTCC |
| System IV_shrinking activator 2 | CCACTCACACTCCACTCCCGCTCGCCACTCTCACTTACAAC |
| Processive Shrinking Directive (PSD) | |
| PSD_A | /5ACryd/CCTAAGTTCGCTGTGGCACCTGCACG |
| PSD_R | /5ACryd/CAACGTGCAGGTGCCACAGCGTGG |
| PSD_growth activator 1 | CCACGCTGTGGCACCTGCACGCACCCACGTGCAGGTGCCACAGCG AACTTAATGATTGTGTATAGT |

| | |
|---|---|
| PSD_growth activator 2 | AGTTAAGAGAATGATTGGGTGCGTGCAGGTGCCACAGCGTAAGTT CGCTGTGGCACCTGCACGTTG |
| PSD_shrinking activator 1 | ACTATACACAATCATTAAGTT |
| PSD_shrinking activator 2 | CACCCAATCATTCTCTTAACT |
| Sequence Design Evolution (SDE) | |
| SDE_A | /5ACryd/CCTAAGTTCGCTGTGGCACCTGCACG |
| SDE_R | /5ACryd/CAACGTGCAGGTGCCACAGCGTGG |
| SDE_v1 (6/3 nT, 0nT spacer, 7nT reversible domain) | |
| SDE_v1_growth activator 1 | CCACGCTGTGGCACCTGCACGCACCCACGTGCAGGTGCCACAGCG AACTTAATGATTG |
| SDE_v1_growth activator 2 | GAATGATTGGGTGCGTGCAGGTGCCACAGCGTAAGTTCGCTGTGGC ACCTGCACGTTG |
| SDE_v1_shrinking activator 1 | CAATCATTAAGTTCGCTGTGGCACCTGCACG |
| SDE_v1_shrinking activator 2 | CGCTGTGGCACCTGCACGCACCCAATCATTC |
| SDE_v2 (6/3 nT, 0nT spacer, 15nT reversible domain) | |

| | |
|---|---|
| SDE_v2_growth activator 1 | CCACGCTGTGGCACCTGCACGCACCCACGTGCAGGTGCCACAGCGAACTTAATGATTGTGTATAGT |
| SDE_v2_growth activator 2 | AGTTAAGAGAATGATTGGGTGCGTGCAGGTGCCACAGCGTAAGTTCGCTGTGGCACCTGCACGTTG |
| SDE_v2_shrinking activator 1 | ACTATACACAATCATTAAGTTCGCTGTGGCACCTGCACG |
| SDE_v2_shrinking activator 2 | CGCTGTGGCACCTGCACGCACCCAATCATTCTCTTAACT |
| SDE_v3 (6/3 nT, 5nT spacer, 15nT reversible domain) | |
| SDE_v3_growth activator 1 | CCACGCTGTGGCACCTGCACGCACCCAGCCATCGTGCAGGTGCCACAGCGAACTTAATGATTGTGTATAGT |
| SDE_v3_growth activator 2 | AGTTAAGAGAATGATTGGGTGCGTGCAGGTGCCACAGCGGCCATTAAGTTCGCTGTGGCACCTGCACGTTG |
| SDE_v3(v2)_shrinking activator 1 | ACTATACACAATCATTAAGTTCGCTGTGGCACCTGCACG |
| SDE_v3(v2)_shrinking activator 2 | CGCTGTGGCACCTGCACGCACCCAATCATTCTCTTAACT |
| SDE_v4 (8/3 nT, 5nT spacer, 15nT reversible domain) | |
| SDE_v4_growth activator 1 (System | CCACGCTGTGGCACCTGCACGCACCCACAGCCATCGTGCAGGTGCCACAGCGAACTTAGGATGATTGTGTATAGT |

| | |
|---|---|
| I_growth activator 1) | |
| SDE_v4_growth activator 2 (System I_growth activator 1) | AGTTAAGAGAATGATTGTGGGTGCGTGCAGGTGCCACAGCGGCCATCCTAAGTTCGCTGTGGCACCTGCACGTTG |
| SDE_v4_shrinking activator 1 (System I_shrinking activator 1) | ACTATACACAATCATCCTAAGTTCGCTGTGGCACCTGCACG |
| SDE_v4_shrinking activator 2 (System I_shrinking activator 1) | CGCTGTGGCACCTGCACGCACCCACAATCATTCTCTTAACT |
| System II_v1 (6/3nT, 5nT spacer, 15nT reversible domain) | |
| System II_v1_growth activator 1 | AAA GCC TAC CAC TCC GTT GCG GAA CCT CTA AGC GCA ACG GAG TGG TAG GCAGACAGGTAAGGTAAGATAGG |
| System II_v1_growth activator 2 | GAATGATTGGGTGCGTGCAGGTGCCACAGCGTAAGTTCGCTGTGGCACCTGCACGTTG |
| System II_v1_shrinking activator 1 | CCTATCTTACCTTACCTGTCTGCCTACCACTCCGTTGCG |
| System II_v1_shrinking activator 2 | GCCTACCACTCCGTTGCGGAACCTCACATCACACTACCC |

| | |
|---|---|
| Trial System T1 | |
| T1_A | /5ACryd/GGAACTCGGCAGTCGTCCAAGCGA |
| T1_R | /5ACryd/ATCTCGCTTGGACGACTGCCGTAT |
| T1_v1_growth activator 1 | ATACGGCAGTCGTCCAAGCGATACGGCTACACTCGCTTGGACGACTGCCGAGTTCCGAGATGTTAGTTAGT |
| T1_v1_growth activator 2 | TGTTGAGTTGAGTTTGCCGTATCGCTTGGACGACTGCCGTACACGGAACTCGGCAGTCGTCCAAGCGAATC |
| T1_v2_growth activator 1 | ATACGGCAGTCGTCCAAGCGATACGGCTCAACTCGCTTGGACGACTGCCGAGTTCCTGATGTTGAGTTGAG |
| T1_v2_growth activator 2 | AGTAGTTTGAATGTTGCCGTATCGCTTGGACGACTGCCGTCAACGGAACTCGGCAGTCGTCCAAGCGAATC |
| T1_v3_growth activator 1 | ATACGGCAGTCGTCCAAGCGATACGGCTACACTCGCTTGGACGACTGCCGAGTTCCGAGATGAGTGTGATA |
| T1_v3_growth activator 2 | GTAGTTGAAGATAGAGCCGTATCGCTTGGACGACTGCCGTACACGGAACTCGGCAGTCGTCCAAGCGAATC |
| Trial System T2 | |
| T2_A | /5ACryd/ATCGGACCAGCACTTCGCCTACGG |
| T2_R | /5ACryd/TGACCGTAGGCGAAGTGCTGGATG |
| T2_v1_growth activator 1 | CATCCAGCACTTCGCCTACGGCTCTACATACACCGTAGGCGAAGTGCTGGTCCGATTGTAGTTAGTTTGAG |
| T2_v1_growth activator 2 | GGAGTGTGAGTGAGTGTAGAGCCGTAGGCGAAGTGCTGGATACAATCGGACCAGCACTTCGCCTACGGTGA |
| T2_v2_growth activator 1 | CATCCAGCACTTCGCCTACGGCTCTACCTAACCCGTAGGCGAAGTGCTGGTCCGATTGTAGTTAGTTTGAG |
| T2_v2_growth activator 2 | TTGAGGGTGAGTTGAGTAGAGCCGTAGGCGAAGTGCTGGCTAACATCGGACCAGCACTTCGCCTACGGTGA |
| T2_v3_growth activator 1 | CATCCAGCACTTCGCCTACGGCTCTACCATACCCGTAGGCGAAGTGCTGGTCCGATGTTGAGTGTTAGATG |

| | |
|---|---|
| T2_v3_growth activator 2 | GGTTGAGAGTGAAGTGTAGAGCCGTAGGCGAAGTGCTGGCATACATCGGACCAGCACTTCGCCTACGGTGA |
| T2_v4_growth activator 1 | CATCCAGCACTTCGCCTACGGCTCTACTTTTTCCGTAGGCGAAGTGCTGGTCCGATTGTGAGTGTAGAGTG |
| T2_v4_growth activator 2 | AGGTGTAGAGTGAGAGTAGAGCCGTAGGCGAAGTGCTGGTTTTTATCGGACCAGCACTTCGCCTACGGTGA |
| Heat-induced reversibility strands | |
| Heat_A | /5Acryd/CTGTCTGCCTACCACTCCGTTGCG |
| Heat_R | /5Acryd/ATTCGCAACGGAGTGGTAGGCTTT |
| Heat_growth activator 1 | AAAGCCTACCACTCCGTTGCGGAACCTCGCAACGGAGTGGTAGGCAGACAG |
| Heat_growth activator 2 | AGGTTCCGCAACGGAGTGGTAGGCCTGTCTGCCTACCACTCCGTTGCGAAT |
| Enzymatic degradation strands | |
| DNase_A | /5ACryd/CCTAAGTTCGCTGTGGCACCTGCACG |
| DNase_R | /5ACryd/CAACGTGCAGGTGCCACAGCGTGG |
| DNase_growth activator 1 | CCACGCTGTGGCACCTGCACGCACCCACGTGCAGGTGCCACAGCGAACTTA |
| DNase_growth activator 2 | TGGGTGCGTGCAGGTGCCACAGCGTAAGTTCGCTGTGGCACCTGCACGTTG |

**Movie S1. Time-lapse fluorescence video of a PAAM-co-BIS-DNA gel over three cycles of DNA-directed growth and shrinking.** Gel size: 1 mm x 1 mm x 60 µm. During each actuation cycle, the gel was first treated with 60 µM System I growth activators containing 1% growth terminators in TAEM-Tween20 buffer. After 72 hours of growth, the DNA solution was removed, 100 µL TAEM-Tween20 was added for 15 mins and removed, and then 150 µL of 60 µM shrinking activator DNA sequences mixed with TAEM-Tween20 was added to induce shrinking. After 24 hours of shrinking, the DNA solution was removed, and 100 µL TAEM-Tween20 was added for 15 mins and removed to prepare for the next actuation cycle. The actuation was recorded using a Syngene gel imager and blue light transilluminator as stated in Methods. Individual frames were subjected to flat fielding and contrast stretching in MATLAB.

**Movie S2-4: Time-lapse fluorescence video of curvature changes of PEG-co-DNA System I-System II bilayer gels during the different actuation programs shown in Fig. 2B-D, respectively.** Gel size: 3.2 mm x 0.8 mm x 0.2 mm at the start of actuation. The actuation was recorded using a Syngene gel imager and blue light transilluminator, as described in the Methods section. Individual frames were subjected to contrast stretching in MATLAB.

**Movie S5: Time-lapse fluorescence video of curvature changes of PEG-co-DNA System III-System IV bilayer gel during the actuation program shown in Fig. 2E.** Gel size: 3.2 mm x 0.8 mm x 0.2 mm at the start of actuation. The actuation was recorded using a Syngene gel imager and blue light transilluminator as stated in Methods. Individual frames were subjected to contrast stretching in MATLAB.

**Movie S6: Time-lapse fluorescence video of DNA-guided letter gel automaton actuation.** Actuation programs are shown in Fig. 3. Gel size: 6 mm x 0.8 mm x 0.2 mm at the start of actuation. The actuation was recorded using a Syngene gel imager and blue light transilluminator as stated in Methods. Individual frames were subjected to intensity processing in MATLAB as stated in Methods.
